# Membrane Proteins at Scale: Automated Copolymer Nanodisc Purification for Structure and Function

**DOI:** 10.1101/2025.09.05.674548

**Authors:** Philipp T. Hanisch, Lea M. Esser, Felipe Merino, Nishika Sabharwal, Kaja A. Reiffert, Sergej Balanda, Lukas Klemm, Ramona V. Busch, Michael Erkelenz, Ana B. Gallego-Bellon, Daniel Erkelenz, Nina V. Heckmann, Christoph Meisen, Oliver Kortheuer, Roland Fabis, Barbara Maertens, Jan Kubicek

## Abstract

Membrane proteins remain among the most important yet least accessible classes of drug targets. Conventional detergents can remove native lipids, destabilizing proteins and limiting downstream biochemistry and structural biology. Amphiphilic copolymers offer a powerful alternative, directly extracting membrane proteins in their native lipid environment, but solubilization outcomes remain unpredictable, turning each new target into a slow empirical search. Here, we introduce an automated, plate-based copolymer screening platform that compresses this process from days to hours using millilitre-scale volumes. Lyophilized copolymer libraries combined with magnetic-bead affinity purification enable parallel testing of dozens of copolymers against multiple targets; across 14 diverse human membrane proteins, next-generation copolymers (AASTY, CyclAPol and Cubipol) systematically outperform classical scaffolds. In this work, we show that this automated copolymer screening robustly identifies the right target-copolymer combination, yielding native-like, active, ligand-binding competent protein suitable for structure determination, and establishes a scalable route to systematic exploration of the membrane proteome.

## Introduction

Membrane proteins (MPs) orchestrate critical processes such as signaling, metabolite transport, and intercellular communication. They constitute roughly one-quarter of the human genome, yet over half of all approved small-molecule drugs act on MPs ^1,2^. Despite this importance, MPs account for less than 5% of entries^3^ in the Protein Data Bank (PDB), underscoring persistent challenges in producing MPs for structural and functional studies.

Over the last decades, detergents have been used to solubilize MPs by mimicking their lipidic environment and shielding their transmembrane domains from the aqueous environment. However, detergent micelles are generally a poor replacement for the native lipid environment. Delipidation can destabilize proteins by disrupting essential lipid-protein interactions, often leading to reduced stability and loss of function ^4–11^. To mitigate this, lipid nanodiscs were introduced, where a membrane scaffolding protein (MSP) derived from apolipoproteins hold a small discoidal patch of membrane ^12^. MSP nanodiscs provide a well-defined phospholipid bilayer. However, MPs must be initially purified in detergents before their reconstitution into MSPs nanodiscs with artificial lipids. By contrast, amphiphilic copolymers such as styrene– maleic acid (SMA) enable direct detergent-free extraction of MPs together with a surrounding belt of endogenous lipids, yielding so-called *native nanodiscs* ^13,14^. These copolymer nanodiscs preserve the native lipid environment around the extracted MP and have been shown to stabilize sensitive lipid-protein interactions ^15,16^.

Building on these foundations, next-generation copolymers have been introduced to improve stability and versatility of these nanodiscs. Examples include AASTY (poly (acrylic acid-co-styrene)) and CyclAPol (Ultrasolute™ Amphipol), which enable broader buffer compatibility combined with high extraction efficiency ^17,18^. The most recent addition is Cubipol, a backbone engineered with multiple chemical modifications to maximize solubilization efficiency, stability, and adaptability across conditions for several downstream applications.

The amphiphilic nature of copolymers itself enables lipid bilayer penetration, nanodisc formation and finally nanodisc stabilization ^19^. SMAs and AASTYs, with aromatic styrene rings, tightly interact with lipid acyl chains, while Ultrasolute^TM^ Amphipols and Cubipols use aliphatic groups for gentler, yet efficient and homogeneous solubilization.

Side chain chemistry is a decisive factor for nanodisc stability and protein compatibility. In unmodified SMA, diisobutylene–maleic acid copolymers (DIBMA) ^20^ and AASTY, the negative charged carboxyl groups ensure solubility and drive nanodisc stabilization by electrostatic repulsion. However, the large number of carboxylic groups makes the copolymers pH sensitive and leads to the unwanted chelation of divalent cations, which restricts their usability in complex or harsh buffer systems. To overcome these limitations, a range of chemical modifications have been introduced.

Glyco-functionalization mimics the natural glycocalyx and improves the compatibility of nanodiscs with glycoproteins, boosting both protein stability and extraction efficiency ^21^. Glycerol-derived groups, such as ethanolamine-modified SMA, enhance hydrophilicity and hydrogen bonding, resulting in tighter size control and improved pH stability, properties that are particularly valuable for NMR studies of small lipid-protein particles ^22^. Quaternary amines, by contrast, introduce cationic or zwitterionic character, which can protect nanodiscs under acidic conditions and in the presence of divalent cations. PEGylation provides steric shielding that suppresses non-specific interactions with ligands or peptides, a critical advantage for GPCR pharmacology and ligand-binding assays ^23^. Finally, zwitterionic sulfobetaine modifications yield more electroneutral nanodiscs that remain stable across wide pH and ionic strength ranges, while also preserving delicate protein-protein and protein-lipid interactions. These properties make them especially suitable for proteomics, mass spectrometry, and charge-sensitive biophysical assays ^24,25^. Such functionalizations were historically distributed across several backbones. Cubipols, however, uniquely condense these diverse chemistries into a single, high-performing backbone with a hydrophobic domain well-suited for different membranes. This integration enables systematic tuning of nanodiscs to the requirements of different proteins.

A key barrier, however, persists: solubilization outcomes and polymer preferences cannot be predicted from the protein’s sequence ^26^. The lipids surrounding MPs, the MP interaction with copolymers as well as the compatibility of the targeted downstream application strongly varies, which makes labor- and time-consuming empirical screening for the right copolymer indispensable ^27,28^.

To address this challenge, we established a universal and semi-automated high-throughput screening platform for copolymer-MP complexes. Our platform minimizes the time and material required for screening, allowing the screening of dozens of different MPs a day. Using 14 diverse MPs, we directly compare SMA, DIBMA, AASTY, Ultrasolute™ Amphipol, and Cubipol, providing the first systematic evaluation of solubilization and stabilization efficiency across multiple MP classes.

As a case study, we focus on the full-length human P2X4 receptor, a clinically relevant ion channel implicated in neuropathic pain and inflammation ^29,30^. We demonstrate that P2X4 can be stabilized in copolymer nanodiscs, remains amenable to electron cryo-microscopy (Cryo-EM), and retains ligand-binding activity. The introduced semi-automated platform transforms empirical copolymer screening into a reproducible and scalable pipeline. It broadens access to functional MPs for structural biology, functional assays, and drug discovery, opening opportunities for previously intractable targets.

## Results

### A universal plate-based robotic system for the rapid and automated purification of membrane proteins in near-native conditions

We have previously shown that the copolymer which solubilizes a specific protein best is not directly predictable from the protein sequence or structure _26_, which makes screening a prerequisite at the start of each new project. Because each copolymer imparts distinct physicochemical properties to the nanodisc environment, systematic screening across different backbones and side-chain chemistries is essential to identify the optimal match for a given MP. The major bottleneck in copolymer-based nanodisc applications is therefore the identification of the right copolymer for each protein target. To address this challenge, we recently introduced our NativeMP™ copolymer suite, comprising 32 copolymers (see Materials and Methods for the complete list of copolymers) that collectively cover most published backbone and side-chain chemistries. Classical native-nanodisc solubilization protocols, such as those originally developed for SMA copolymers, required ∼20 hours of manual work to purify a single protein in one copolymer, making broad screening impractical (Fig. 1a). Our optimized screening protocol for this copolymer suite reduces the manual screening time to ∼6.5 hours (see Material and Methods). The process includes a magnetic-bead-based purification step, aimed at determining not only the best copolymer for solubilization, but one that provides a pure, monodisperse protein afterwards. Despite the throughput improvement, the full protocol remains both highly resource and labor intensive. Considering the nature of the screening, we reasoned that the full protocol could be automated using a liquid handler/magnetic bead transfer instrument such as a KingFisher Flex. As such, we sought to create a fully robotized version of our copolymer suite, aiming to reduce processing time as well as the material required. To this end, we developed a protocol requiring two types of 96-well plates. One of the plates contains the set of lyophilized, prebuffered copolymers to be screened against, referred as the NativeMP^TM^ copolymer plate. The second contains dehydrated magnetic beads (MagBeads) carrying an affinity purification system, referred as PureHT^TM^ plate. The beads can be quickly resuspended by mechanical mixing, with the process having no significant impact on binding capacity. For membrane proteins, we commonly use the C-terminal Rho-tag and beads with Rho 1D4 antibody ^31^. Not only is the Rho 1D4 antibody compatible with native nanodiscs, but its very high affinity for the Rho-Tag is key when working with proteins expressed at low levels. However, other purification strategies, such as Step-tag**^®^**/Strep-Tactin**^®^**XT, His-Tag/Ni-INDIGO as well as Ni-NTA or FLAG^®^-Tag, are fully compatible with our protocol. Fig 1b shows a schematic representation of the automated purification procedure. Briefly, after cell disruption via sonication, the supernatant of a 9,000 x g centrifugation is directly loaded onto a NativeMP^TM^ copolymer plate for automated solubilization (Fig. 1c). After 5 min time, MagBeads are transferred from a Rho PureHT^TM^ plate to the solubilized membranes for binding. The samples are thoroughly washed, and proteins are then eluted using free Rho peptide (see methods for a detailed protocol). Using a KingFisher Flex purification system, the complete procedure from cell pellet to eluates requires 90 min (Fig. 1c). Moreover, using a 96-well plate format, each well takes 1000 µL of lysate, reducing the amount of eucaryotic cell culture suspension needed for a full screening to around 300 mL for a full 32 set copolymer screen.

**Figure 1.**
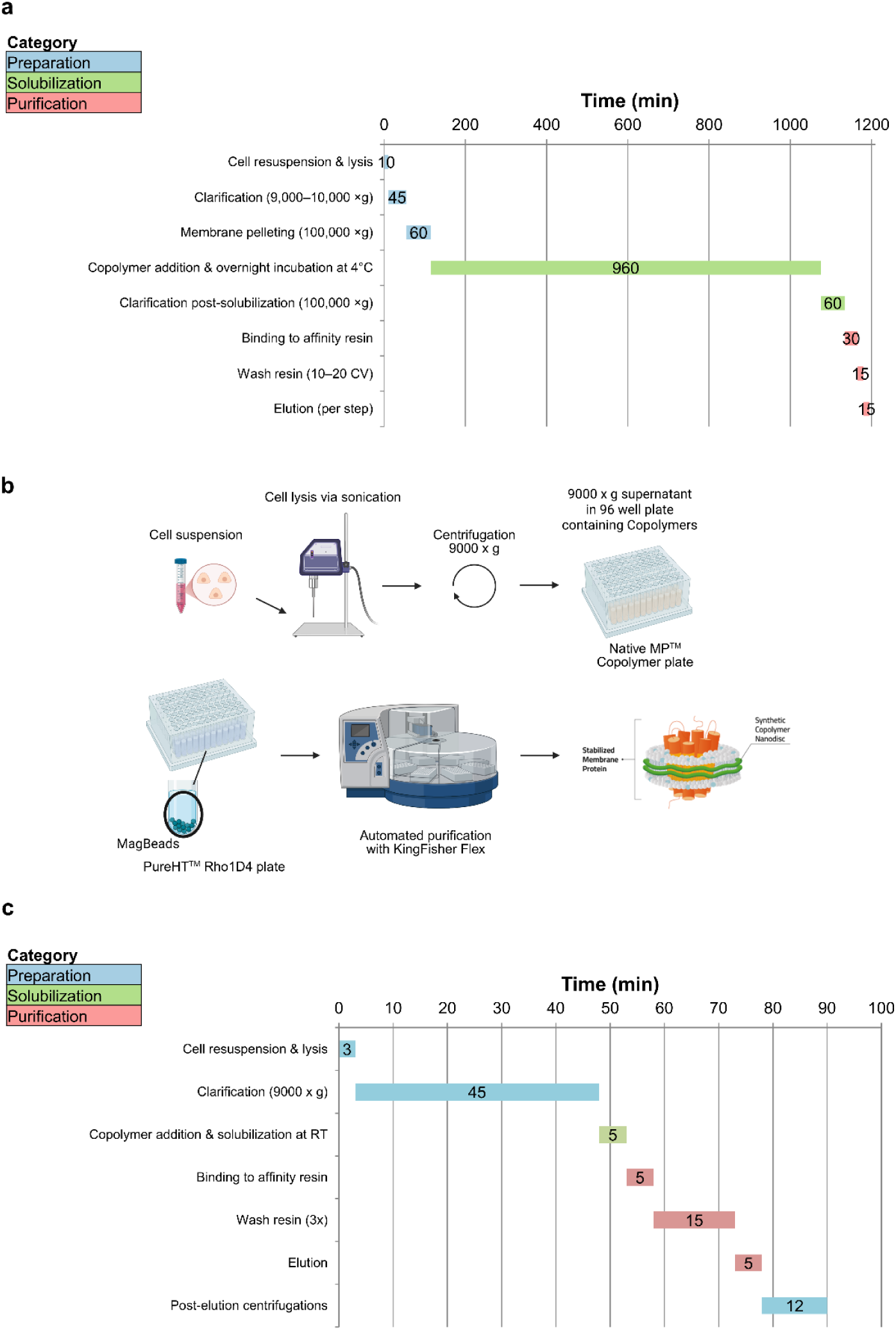
Comparison of classical and automated copolymer workflow. a. Classical manual workflow as described in the literature, comprising cell lysis, clarification, membrane pelleting, copolymer-mediated solubilization, secondary clarification, and affinity purification. The full procedure requires ∼1200 min. Preparative, solubilization, and purification steps are indicated in blue, green, and red, respectively. b. Schematic overview of the adapted semi-automated workflow using pre-buffered NativeMP™ copolymer plates and PureHT™ magnetic bead plates on a KingFisher Flex system. After lysis and clarification, clarified lysates are directly solubilized and purified without membrane pelleting. c. Gantt chart illustrating the streamlined workflow, reducing the total processing time to ∼90 min (7.5% of the manual protocol). Eluates from the semi-automated workflow are of high purity and directly compatible with downstream biophysical analyses (DLS, nDSF), classical assays (SDS–PAGE, Western blot), and Cryo-EM workflows.

### Plate-based copolymer screening reveals broad applicability across membrane protein families

To test whether our plate-based system could be used as a universal tool for copolymer screening of MP targets, we screened 14 different Rho-tagged human membrane proteins with 15 selected copolymers from all six backbones (see Materials and Methods for the complete list of copolymers). We chose a diverse set of full-length, wildtype MPs, including GPCRs (HTR7, ADBR2, AMYR3), solute carrier transporters (SLC1A6, SLC2A1, SLC15A4), voltage-gated channels (hERG^T^), claudins (CLDN1, CLDN3, CLDN4), and other cell surface proteins (CD68, CD163, EGFR, FAS). Many of these proteins, particularly GPCRs, are considered extremely hard targets to work with, particularly due to very poor expression and solubilization efficiency with classical methods combined with low stability. Fig. 2a shows a typical result for a GPCR: the serotonin receptor 7 (HTR7).

**Figure 2.**
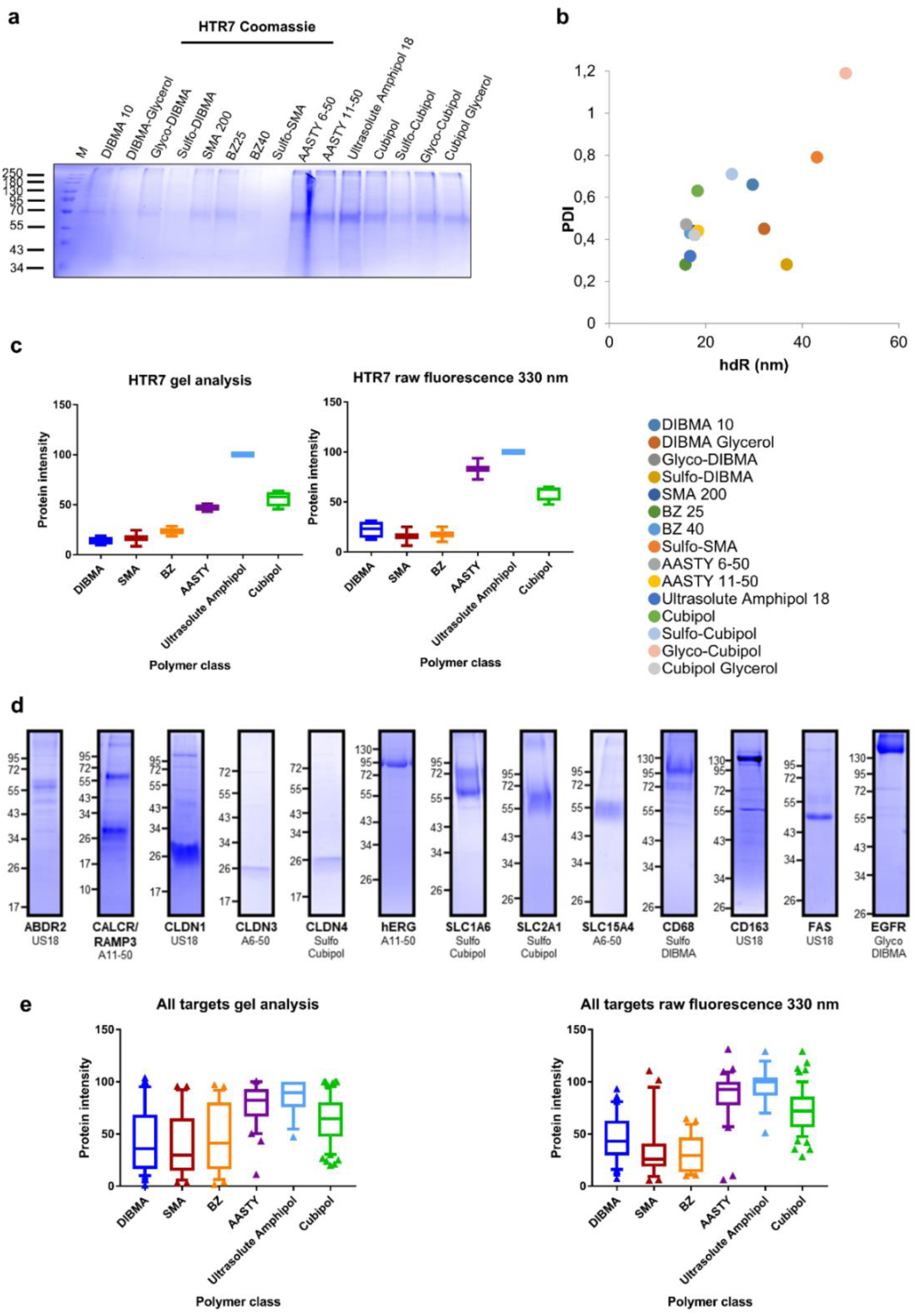
Quantitative analysis of membrane protein content and quality by SDS–PAGE, 330 nm fluorescence and DLS. a., d. Solubilization and purification efficiency was screened for 14 MPs with 15 different NativeMP^TM^ copolymers. Eluates were run an SDS–PAGEs (Coomassie stained). c., e. Protein quantity was analyzed either by densitometry (ImageJ Gel Analyzer) or by buried tryptophan fluorescence at 330 nm (NanoTemper Panta Discovery). Intensities were normalized to the strongest signal in each dataset (set to 100%) and plotted as box plots to visualize distributions across copolymer families. d. For HTR7, eluates from all tested copolymers are shown (a), while for the remaining targets only the strongest eluates are displayed. Full, unmodified SDS–PAGEs and Western blots are provided in Fig. S1-3. In panel e, results from all 14 targets were combined to compare solubilization and stabilization efficiencies across the six copolymer backbones. Panel b shows DLS of HTR7 eluates, with hydrodynamic radius (x-axis) plotted against PDI (y-axis).

With most copolymers, SDS-PAGE shows a single band corresponding to HTR7. Quantification of the HTR7’s band intensity clearly indicates that next-generation copolymers (AASTYs, Ultrasolute^TM^ Amphipols, and Cubipols) outperform their classical counterparts in solubilization power (Fig. 2c). A similar conclusion can be drawn from the total fluorescence measured at 330 nm (measuring the nonpolar/buried environment of tryptophan residues) which correlates with the amount of purified protein (Fig. 2c). Dynamic light scattering (DLS) analysis identified six copolymers that yielded particles within a hydrodynamic radius <20 nm (Fig. 2b, Extended Data Fig. 5b). Among these, Ultrasolute^TM^ Amphipol 18 (U18), AASTY 11-50 and three Cubipol variants stood out for combining smaller particle size with low polydispersity, while the AASTY 6-50 and BZ40 produced particles close to the size threshold (Extended Data Fig. 5b). By contrast, several copolymers produced larger or more heterogeneous assemblies.

Overall, while a subset of copolymers reliably produced nanodiscs below 20 nm with acceptable monodispersity (>0.4), Sulfo-SMA, DIBMA Glycerol and Sulfo DIBMA tended to yield larger, more polydisperse assemblies, highlighting the strong influence of backbone and side-chain chemistry on nanodisc homogeneity (Fig. 2, Extended Data Fig. 5b). Similar variations in hydrodynamic radii have also been described before for other targets ^32^.

### Systematic evaluation reveals reproducible and superior solubilization efficiencies of next-generation copolymers

These results are not unique to HTR7. Across all tested membrane protein targets, we obtained eluates of high purity for most copolymer-protein complexes, albeit with varying solubilization efficiency (Fig. 2 d,e; Extended Data Fig. 1-3). As expected, the optimal copolymer differed between targets. Nonetheless, statistical analysis across the six copolymer backbones and the 14 analyzed targets revealed clear trends: classical SMA and DIBMA displayed broad yield distributions with lower relative median yields (29.7% and 36.0% in Coomassie; 25.8% and 43.0% in 330 nm fluorescence), whereas next-generation copolymers systematically performed better. AASTYs and Ultrasolute^TM^ Amphipols achieved the highest medians (82.4% and 89.7% in Coomassie; 92.6% and 100% in 330 nm fluorescence), while Cubipols combined strong median values (64.7% and 72.1%) with narrow variance, indicating high reproducibility across targets (Fig. 2e, Extended Data Fig. 4).

SMA remains the most widely applied copolymer in the field, representing the current “gold standard” for native nanodiscs. Statistical analysis of our gel quantifications shows that AASTY (p < 0.0001), Ultrasolute^TM^ Amphipol (p < 0.0001) and Cubipol (p = 0.0002) outperform SMAs in solubilization yields. Conversely, DIBMA (p = 0.479) or SMA BZ (p = 0.430) are statistically indistinguishable classical SMAs (see methods and Extended Data Fig. 4 for details). A corresponding statistical evaluation of the fluorescence at 330 nm corroborates this result, underscoring the consistency of the performance differences across independent readouts (Extended Data Fig. 4b). Thus, the performance gap between classical and next-generation copolymers is not only systematic but also statistically robust, with next-generation copolymers consistently providing superior yields.

### Target-by-target DLS analysis highlights next-generation copolymers as the most reproducible performers

To assess the generalizability of copolymer performance, we correlated the hydrodynamic radius (hdR) and the polydispersity index (PDI) values for each target-copolymer combination of all 14 screened targets (Extended Data Figure 5. b). The majority of next-generation copolymers produced nanodiscs with hydrodynamic radii below 20 nm, and PDIs below 0.4 indicating high monodispersity, while several classical copolymers displayed broader distributions.

As the optimal copolymer depends on the target MP, we ranked all copolymers by lowest hdR in combination with lowest PDI and extracted the three best-performing conditions (Extended Data Fig. 5a). This analysis revealed that, although the precise ranking differed between proteins, Cubipols, AASTYs, and Ultrasolute^TM^ Amphipols were mostly within the Top-3 across nearly all targets. For example, U18 appeared among the Top-3 in 7 of 14 targets, Cubipol derivatives in 13 of 14 targets, and AASTY variants in 5 of 14 targets. Classical SMA and DIBMA copolymers reached the Top 3 in 5 and 2 of 14 targets: SMA 200 and Glyco-DIBMA were the only classical representatives that occasionally yielded acceptable size and dispersity values. This correlates with the gel analysis (Fig. 2d, Extended Data Fig. 1-3) where U18 showed highest output in 9 out of the 14 targets, Cubipol derivates in 2 and AASTY variants in 3.

Taken together, this demonstrates that while nearly all copolymers can solubilize MPs, systematic screening is essential to pinpoint the optimal match for each target. The Top-3 analysis further illustrates that next-generation copolymers consistently outperform first-generation backbones, delivering smaller and more homogeneous nanodiscs that are directly suitable for downstream structural and biophysical workflows.

### Comparative copolymer screening of P2X4^T.ni^ and P2X4^HEK^ highlights expression system differences

While monodisperse and pure protein samples are a prerequisite for downstream analysis, the true strength of copolymer-based nanodiscs lies in their ability to preserve the native lipid environment that underpins correct folding and function. To directly demonstrate this advantage, we next turned to P2X4 as a case study. The ion channel P2X4 is a well-characterized 120 kDa trimeric protein and clinically relevant purinergic receptor, with several known ligands^33–35^, making it a perfect candidate for an in-depth biophysical and structural characterization.

We expressed P2X4 in two eukaryotic hosts, *T.ni* insect cells and HEK293 mammalian cells, to decouple sequence-intrinsic effects from the influence of the membrane and post-translational landscape, and to benchmark our screen across divergent, yet widely used, production backgrounds. *T.ni* cells offer high, scalable yields and a simplified N-glycosylation profile that is often advantageous for biochemical handling and certain structural workflows. In this context we chose a *T.ni-cells expression optimized* construct (P2X4^T.ni^) carrying three mutations (N75R, N184R, and E307E) designed to reduce glycosylation and enhance ATP binding. In contrast, we chose the wild-type human P2X4 (P2X4^HEK^) for HEK293 cells. This setting provides the closest approximation to the *in vivo* environment, making it highly suitable for evaluating copolymer nanodiscs in terms of folding, function, and ligand interactions. Using both hosts therefore (i) tests generalizability of copolymer performance across distinct membrane backgrounds, (ii) reveals host-dependent differences in nanodisc size, monodispersity, and thermostability, and (iii) guides practical choices (yield vs. native-likeness) for downstream activity assays and structural determination.

For both expressions systems, we performed a copolymer screening against the full NativeMP^TM^ copolymer suite Cube Biotech provides (see copolymer list in Material and Methods part). As shown with our previous targets next-generation copolymers provide significantly higher protein yields for both P2X4 variants (Fig. 3, Extended Data Fig. 6). Interestingly, the P2X4^HEK^ samples are of higher quality than P2X4^Tni^, both in terms of purity and yield (Fig. 3). Although this could be partly attributed to the small sequence differences between them, the lipidic microenvironment around the protein likely impacts the solubilization capacity of the copolymers, as we have proposed previously ^26^. To gain further insight into the biophysical properties of both constructs, we first analyzed three out of the 32 copolymer stabilized complexes in detail: SMA200, AASTY 6-55 and Glyco-Cubipol (Fig. 4a-d). The *Tni*-derived samples were more heterogeneous, in contrast, the HEK-derived samples were markedly more homogeneous demonstrating compact and monodisperse assemblies (Fig. 4 a,c, Extended Data Fig. 7).

**Figure 3.**
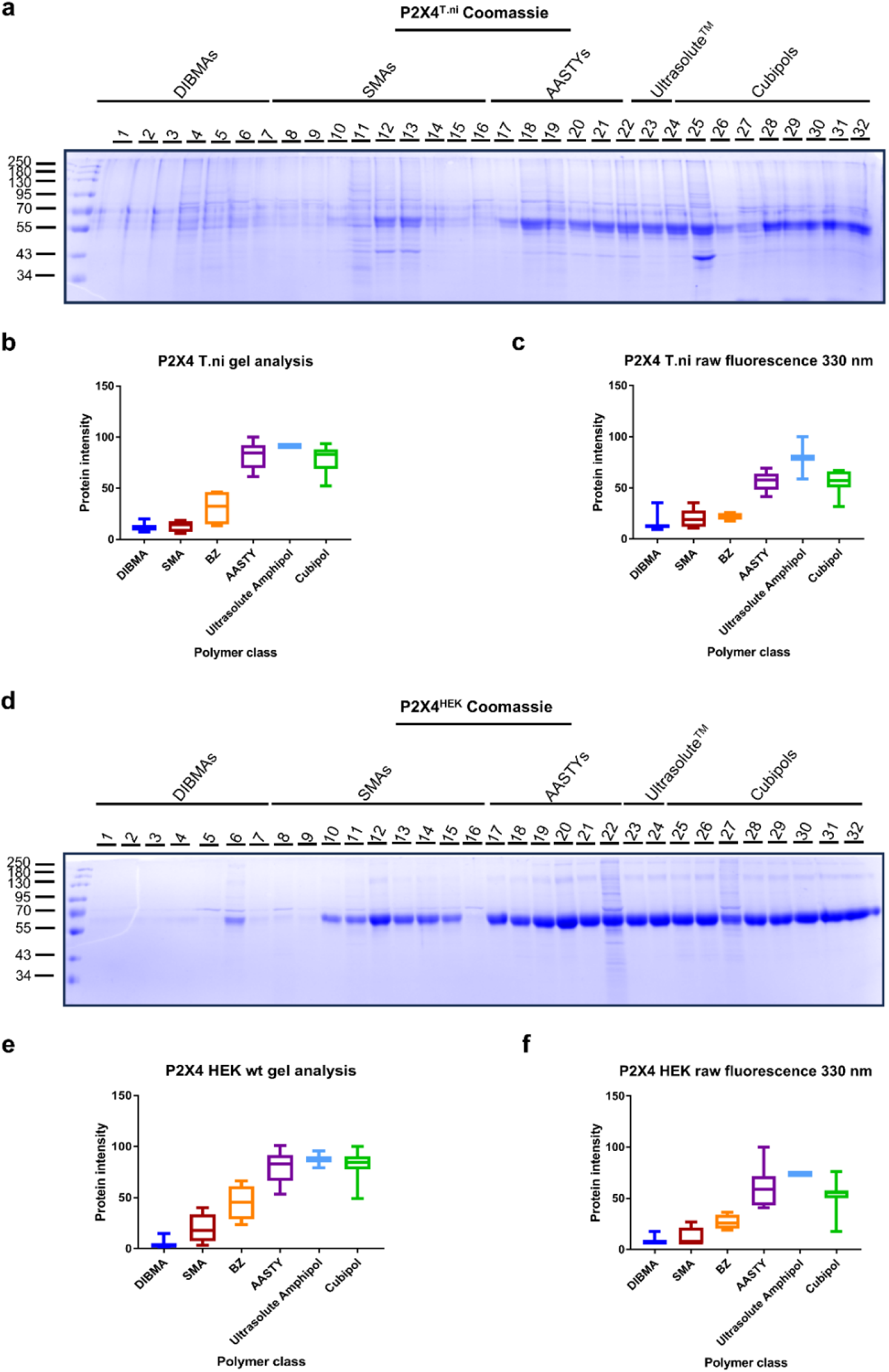
Comparative NativeMP^TM^ 32 copolymer screening of P2X4^Tni^ and P2X4^HEK^. a., d. Solubilization and purification efficiency of 32 different copolymersc (full list in material and methods) was screened for P2X4^Tni^ and P2X4^HEK.^. SDS–PAGEs of the eluates were analyzed either by densitometry (ImageJ Gel Analyzer) (b,e) or by buried tryptophan fluorescence at 330 nm (NanoTemper Panta Discovery) (c,f). Intensities were normalized to the strongest signal in each dataset (set to 100%) and plotted as box plots to visualize distributions across copolymer families. Corresponding WBs can be viewed in Fig. S6.

**Figure 4.**
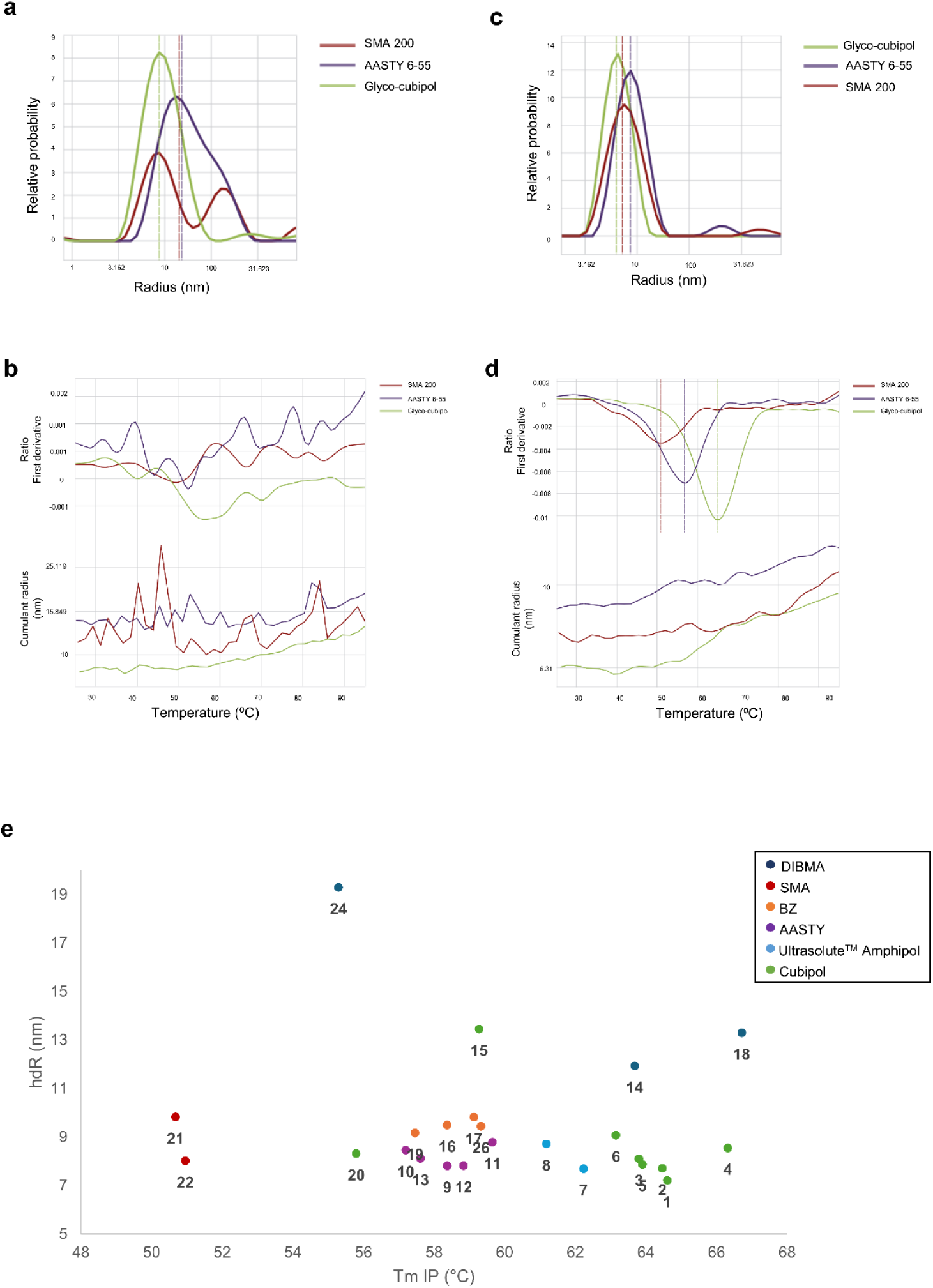
Biophysical analysis and copolymer ranking of P2X4. a. Dynamic light scattering (DLS) of P2X4^Tni^ solubilized in SMA200, AASTY 6-55, and Glyco-Cubipol revealed heterogeneous particle populations with large hydrodynamic radii (hdR) and high PDIs. b. NanoDSF of the same P2X4^Tni^ samples showed no detectable inflection points (IPs), consistent with conformational heterogeneity. c. DLS of P2X4^HEK^ solubilized in SMA200, AASTY 6-55, and Glyco-Cubipol demonstrated compact and homogeneous nanodiscs with hdR values of 7.3 nm (PDI 0.50), 8.5 nm (PDI 0.10), and 7.9 nm (PDI 0.29), respectively. d. NanoDSF of P2X4^HEK^ revealed distinct IPs at 50.9 °C (SMA200), 56.7 °C (AASTY 6-55), and 65 °C (Glyco-Cubipol). e. Copolymer screening of full-length P2X4 expressed in HEK293 cells. Thirty-two copolymers representing six backbones were tested for solubilization and stabilization efficiency. Following purification, eluates were analyzed by Coomassie-stained gels for yield and by biophysical methods including T_m_, hdR, PDI, and turbidity. Values were normalized and integrated into a composite score (T_m_ 0.35, hdR 0.25, PDI 0.10, turbidity 0.05, elution quantity 0.25). Copolymers were ranked by this score, with lower ranks indicating superior performance. The scatter plot displays T_mIP_ (x-axis) versus hdR (y-axis), visualizing stability and size characteristics. Top performers included several Cubipol derivatives (e.g., Glyco-Cubipol, Sulfo-Cubipol Medium, Cubipol PEG, Cubipol Amine; shown in green), which consistently combined high thermostability with favorable size distribution and yield. The full dataset is provided in Figure S9, S7,8.

### Multifactorial evaluation of P2X4-copolymer complexes identifies optimal candidates for functional and structural studies

One of the common challenges when working with eukaryotic membrane proteins is their relatively low stability, which constrains both assay duration and the temperatures at which experiments can be performed. We reasoned that preserving native lipids in copolymer nanodiscs should provide additional stabilization. Indeed, nanoDSF analysis demonstrated that the copolymer environment enhanced P2X4^HEK^ thermostability without requiring additional stabilizing mutations or kosmotropic agents (Fig. 4d). Clear inflection points were detected at 50.9 °C for SMA200, 56.7 °C for AASTY 6-55, and 65 °C for Glyco-Cubipol (Fig. 4d), underlining how the choice of copolymer directly impacts receptor stability. In contrast, no transition temperatures could be detected for any of the *Tni*-cell derived samples (Fig. 4b).

For activity assays and structural determination, we are ultimately looking for target-copolymer combinations that deliver particles with low PDI, small hydrodynamic radii, sufficient thermal stability to allow measurements at room temperature or up to 37 °C, minimal turbidity even under stress conditions, and as much protein as possible in the eluate. We therefore carried out a multifactorial analysis of the P2X4^HEK^ screen and ranked the copolymers accordingly (Fig. S7). The ranking was generated by weighting the different parameters as follows: Tm (0.35), hydrodynamic radius (0.25), polydispersity index (0.10), turbidity (0.05), and elution quantity (0.25). A score of 1 is optimal. Elution quantity was derived from gel densitometry (Fig. 3e) and integrated as a direct measure of protein yield. For clarity, the ranking was visualized in a scatter plot with T_m_ values on the x-axis and hydrodynamic radii (hdR) on the y-axis (Fig. 4e; complete ranked dataset is provided in Extended Data Figure 7). Using this scoring scheme, Cubipol derivatives dominated the top of the ranking, with five Cubipol variants (Glyco-, Sulfo-, PEG-, Amine-, and Glycerol-modified forms) and Cubipol occupying positions 1-6 (Fig. 4e, green, Extended Data Fig. 7). These copolymers consistently produced eluates with high thermal stability, small hydrodynamic radii, and low polydispersity, underlining their ability to stabilize P2X4 in a native-like environment. Immediately following the Cubipols, Ultrasolute^TM^ Amphipols ranked highly (positions 7–8, Fig. 4e, light blue), yielding small and highly homogeneous nanodiscs. AASTYs occupied the mid-field (positions 9–13, Fig. 4e, purple), with reasonable stability but somewhat higher PDI values. By contrast, DIBMAs and classical SMAs clustered at the bottom of the ranking (positions 14 and below, Fig. 4e, dark blue; red), associated with oversized or polydisperse particles and reduced stability. Although some individual representatives performed slightly better than their family averages, overall, they did not reach the level of the next-generation copolymers.

During the screen we observed that in all cases, the protein migrated at a higher apparent molecular weight on SDS-PAGE, consistent with extensive N-glycosylations at multiple asparagine residues (N75, N110, N153, N184, N199, and N208). To directly confirm this for the HEK construct stabilized in biotinylated U18, we performed a PNGase F digestion, which removes N-linked glycans. Following deglycosylation, P2X4^HEK^ shifted to a lower apparent molecular weight, demonstrating that the observed migration pattern is indeed caused by heavy glycosylation rather than differences in the polypeptide backbone (Fig. 5a). For both expression systems, P2X4 could be purified as the expected trimer. For the HEK-derived construct we confirmed the expected trimeric state directly by native PAGE (Fig. 5b), further validating that the receptor remains assembled in its physiological oligomeric form.

**Figure 5.**
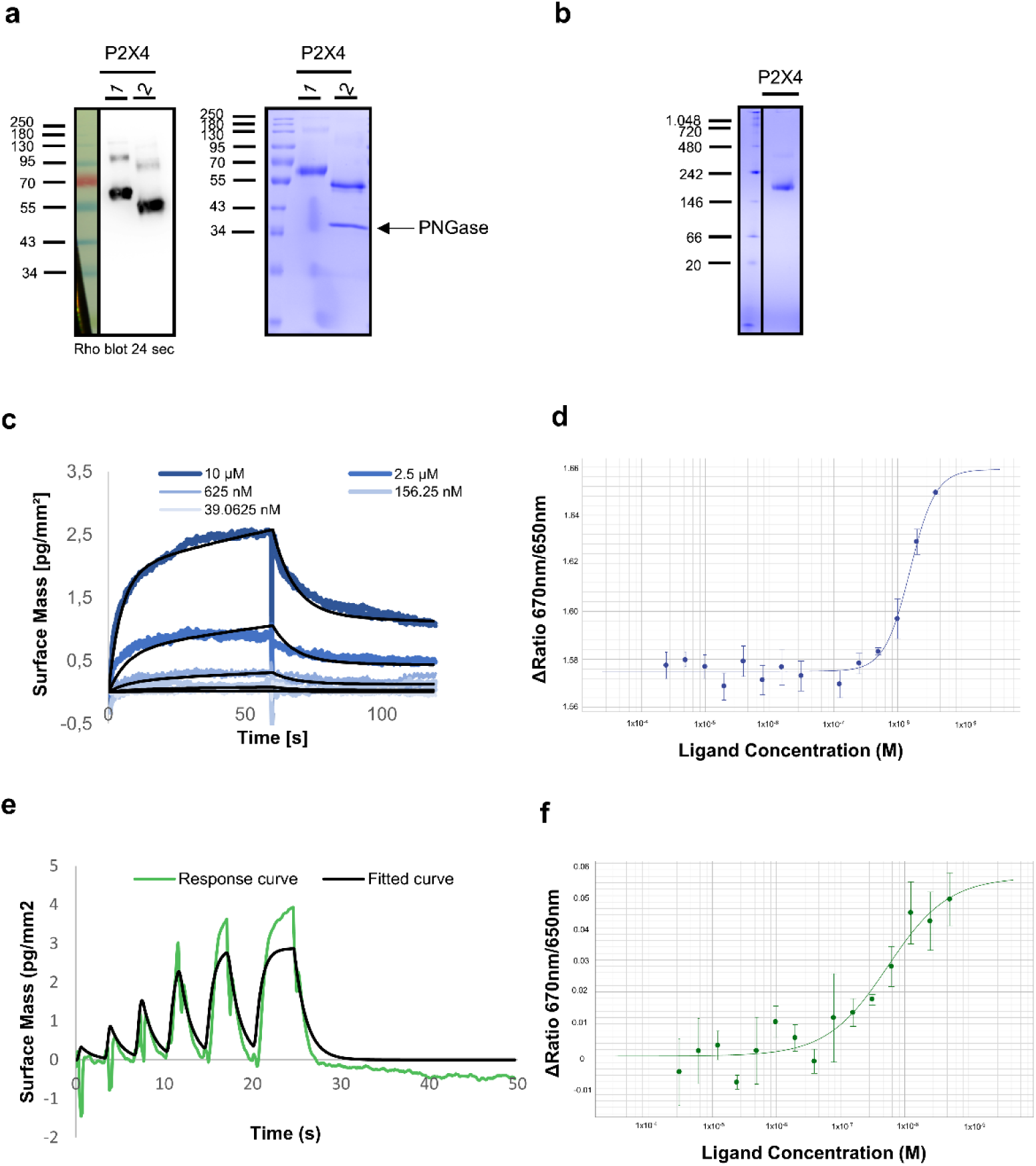
Biochemical characterization and ligand-binding characterization of P2X4^HEK^. **a.** Anti-rho western blot and SDS-PAGE analysis of the PNGase digestion of P2X4^HEK^. A clear shift of the P2X4 bands before (1) and after (2) digestion demonstrates that the higher apparent molecule weight is due to glycosylation of the protein. **b**. Native PAGE showing that P2X4^HEK^ is assembled as a trimer. **c**. Grating-coupled interferometry (GCI) sensorgrams of BAY-1797 binding to P2X4^HEK^ reconstituted in U18 biotin nanodiscs. Serial twofold dilutions (39 nM–20 µM) were injected at 25 °C onto immobilized P2X4–U18 biotin nanodiscs. Sensorgrams were blank/DMSO corrected and globally fitted to a 1:1 Langmuir binding model, yielding a dissociation constant (Kd) of 1.3 µM with kinetic rate constants k_on_= 9.28 ± 3.03 x 10^3^ M^−1^S^−1^ and k_off_ = 1.15 ± 0.30 × 10^−1^ S^−1^ **d.** Spectral shift analysis of BAY-1797 binding to fluorescently labelled full-length P2X4^HEK^ in U18 biotin nanodiscs. Fluorescence shifts (ΔRatio 670/650 nm) were recorded upon titration of BAY-1797 (twofold serial dilution) and fitted by nonlinear regression, yielding a K_d_ of 15.6 µM. **e.** GCI WaveRAPID analysis of 5-BDBD binding to P2X4^HEK^–U18 biotin nanodiscs. Sensorgrams were referenced against empty nanodisc controls and fitted to a conformational-change binding model, giving a K_d_ of 10 µM. **f.** Spectral shift assay of 5-BDBD with P2X4^HEK^ in U18 biotin nanodiscs. Titration of fluorescently labelled receptor with serial dilutions of ligand produced concentration-dependent fluorescence shifts, yielding a binding affinity of 5.4 µM.

### Native nanodiscs allow the determination of ligand-binding parameters in near-native conditions

The P2X4^HEK^ screen clearly identified Cubipols as the top-performing copolymer family in terms of yield, stability, and monodispersity. For the functional validation, however, we selected U18 as a representative next-generation copolymer outside the Cubipol family. This ensured that our results did not rely exclusively on Cubipols and demonstrated that the NativeMP^TM^ approach is broadly applicable across different copolymer classes. U18 proved to be highly stable and reproducible across independent preparations and could be readily scaled up to provide sufficient material for biophysical and biochemical analyses in parallel. In this way, Cubipols set the benchmark for overall performance in the screen, while U18 provided an orthogonal, well-established reference for functional assays.

To validate receptor activity, we used two complementary techniques that probe different aspects of protein-ligand interactions: solution-based spectral shift analysis (NanoTemper Monolith X) (Fig. 5d,f) and surface-based grating-coupled interferometry (Malvern Panalytical GCI) (Fig. 5c,e). For GCI, U18 was chemically modified with biotin, which allowed nanodiscs to be immobilized directly on a streptavidin-coated sensor chip (Extended Data Fig. 10). This provided a stable anchor while still preserving the receptor’s conformational flexibility and free access to its ligand-binding pocket. To account for non-specific interactions between analytes and the copolymer surface as well as the phospholipid environment, we implemented a dedicated reference channel containing control copolymer-host lipid nanodiscs. These nanodiscs were assembled from host-cell lipids and the same copolymer matrix used for protein solubilization, but without incorporated P2X4. This design ensured that both reference and sample surfaces shared identical physicochemical properties, allowing the specific receptor-ligand response to be isolated from background signals arising from the copolymer or lipid environment. The inclusion of such empty copolymer nanodiscs as reference channels represents an important methodological advance for copolymer-based biosensing platforms including GCI, SPR, and BLI by enabling quantitative correction of non-specific binding contributions and more accurate determination of kinetic and affinity parameters. To ensure consistency across methods, we used the same biotinylated U18 nanodiscs for spectral shift analysis, where P2X4 was fluorescently labelled via the NanoTemper Spectral Shift Optimized Protein Labeling Kit (NT-L021). This setup enabled us to directly monitor ligand-induced spectral changes in solution and compare the results with the surface-based measurements under identical biochemical conditions.

Afterwards the receptors interaction with two selective antagonists, 5-BDBD and BAY-1797, was tested. For 5-BDBD, spectral shift produced an IC₅₀ of 5.4 µM (Fig. 5f), while GCI measured a dissociation constant (Kd) of 10 µM (Fig. 5e). These results align with previously reported EC₅₀ values in the 0.5–5 µM range ^36^, confirming that P2X4 retains its pharmacological properties in biotinylated U18 nanodiscs. For BAY-1797, both techniques also detected binding. Spectral shift gave a Kd of 15.6 µM (Fig. 5d), while GCI reported a stronger interaction at 1.3 µM (Fig. 5c). These results are demonstrating that biotinylated U18 stabilization preserves both the structure and function of P2X4. The biotinylated U18 nanodiscs not only kept the receptor stable across assay types but also provided the flexibility needed to cross-validate results under different experimental conditions. Consistent binding parameters were obtained in both solution and immobilized formats presses on the robustness of the NativeMP^TM^ technology.

### The NativeMP^TM^ platform is directly compatible with Cryo-EM structural determination

Our biophysical characterization demonstrates that copolymer-solubilized P2X4^HEK^ is trimeric, glycosylated, and can bind its known ligands with the expected affinity. Next, we sought to demonstrate that the NativeMP^TM^ platform can preserve the native structure of the target protein. To this end, we decided to use Cryo-EM, as recent advances have positioned it as the go-to technique for the structural determination of membrane proteins ^37^. The P2X4^HEK^ screening resulted in small volumes of samples with exceptionally high biochemical quality (Fig. 3d). Although this might limit some downstream applications, we wondered whether they could be directly used for Cryo-EM screening. Remarkably, eluates taken directly from the 96-well screening plate produced grids with well-distributed P2X4 particles, showing the characteristic triangular shape of the protein (Fig. 6a). To further test these samples, we collected small data sets for Cubipol Glycerol and Sulfo-Cubipol medium. 3D refinement of the Glycerol-Cubipol data produced a reconstruction of the protein at 3.6 Å resolution (Fig. 6b,c Extended Data Fig. 11a, Table S1). Notably, clear density can be observed for an ATP molecule at the agonist-binding site, which was likely captured by the protein upon cells disruption (Fig. 6c). This suggests that the receptor remains functional within the copolymer nanodisc. Interestingly, comparison with a previously determined structure of P2X4 ^38^ shows that while the extracellular domain is in the open conformation, the transmembrane helices are in unique state, different from the desensitized or closed states of the channel (Extended Data Fig. 12).

**Figure 6.**
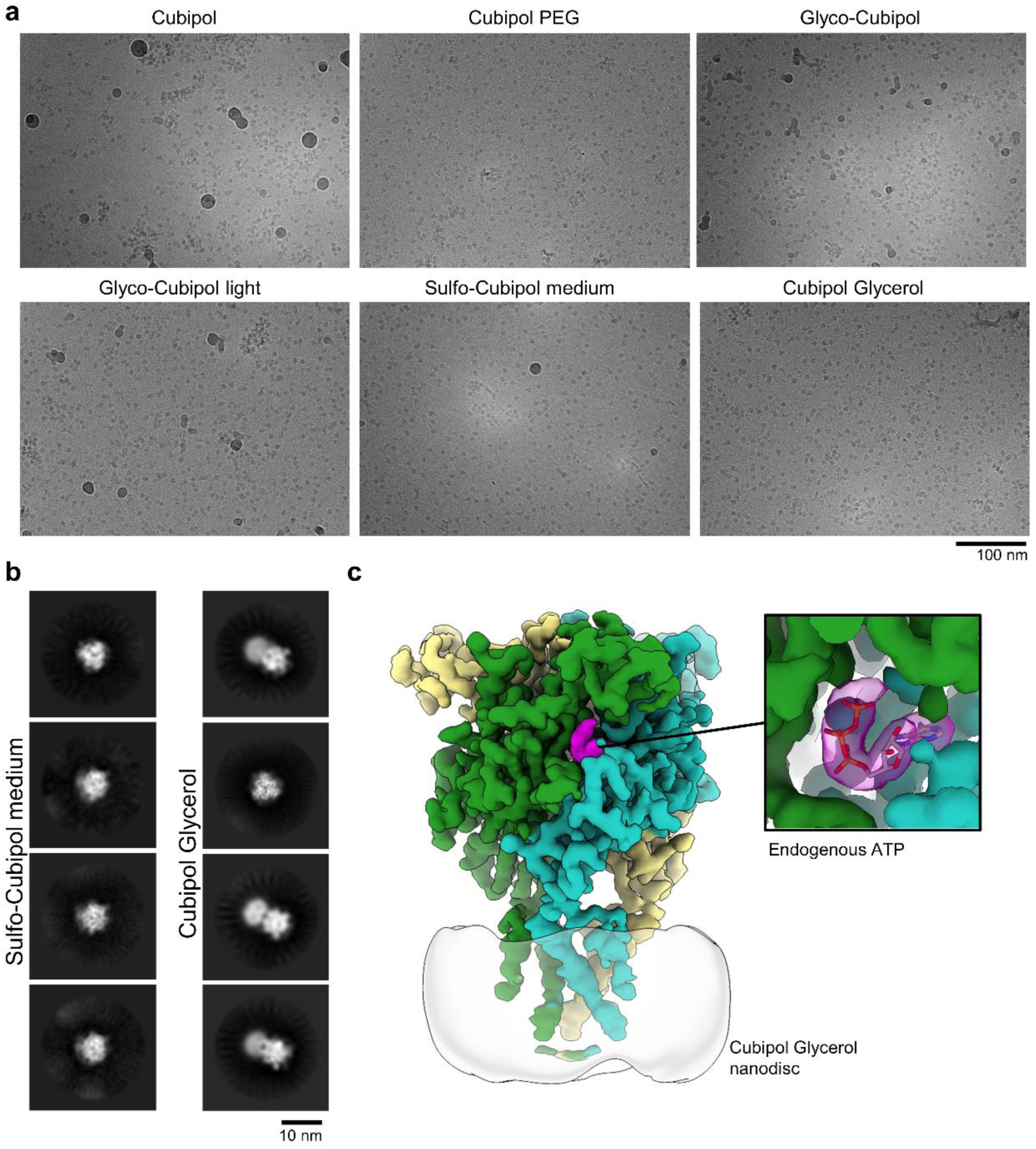
Cryo-EM screening P2X4^HEK^ samples purified with our automated protocol. **a.** Representative micrographs for all tested polymers, collected at ∼ 3 µm defocus. **b**. Representative 2D classes obtained from the small data sets collected for Sulfo-Cubipol medium and Cubipol Glycerol. Sufo-Cubipol medium shows strong preferential orientation, while Cubipol Glycerol has a more balanced orientation distribution. **c**. 3D reconstruction of the Cubipol Glycerol sample. Endogenous ATP co-purified with the sample is highlighted in magenta, and shown in more detail in the close-up panel.

### Cryo-EM optimization of P2X4^Tni^ by targeted copolymer selection and upscaling

To further challenge the platform, we turned to the P2X4^Tni^ construct, which had shown lower stability and higher heterogeneity in our screening (Fig. 3a; Fig. 4a,b). Here, robotic eluates alone were of lower quality, necessitating upscaled purifications with selected copolymers. We focused on Glyco-DIBMA, Ultrasolute^TM^ Amphipol 18, Glyco-Cubipol, Glycerol-Cubipol, and Sulfo-Cubipol (See Extended Data Tab. 1 for microscopy and dataset size details). Well-distributed particles with the expected P2X4 shape were clearly visible in all samples (Extended Data Fig. 11b). In all cases, we obtained 2D classes showing high-resolution features (Fig. 7a). However, all samples show strong orientation bias, with most particles arranged in top-view orientation (Fig. 7a). This was particularly severe for Glyco-DIBMA, where no 3D reconstruction was possible. For the remaining copolymers, we obtained 3D reconstructions with resolutions ranging from ∼7 Å to 2.9 Å (Fig. 7a, b, Extended Data Fig. 11a, Extended Data Tab. 1, Extended Data Tab. 2). We attribute the lower resolutions observed with U18 and Glyco-Cubipol to poor particle signal-to-noise ratio, which can stem from either tick ice or the bulkiness of copolymer side chains. Both of these issues could be mitigated by optimized grid preparation or larger data collection. Indeed, high-resolution structures of U18 solubilized AcrB ^18^ and MsbA ^39^ have been previously reported, demonstrating that high-resolution possible is technically possible.

**Figure 7.**
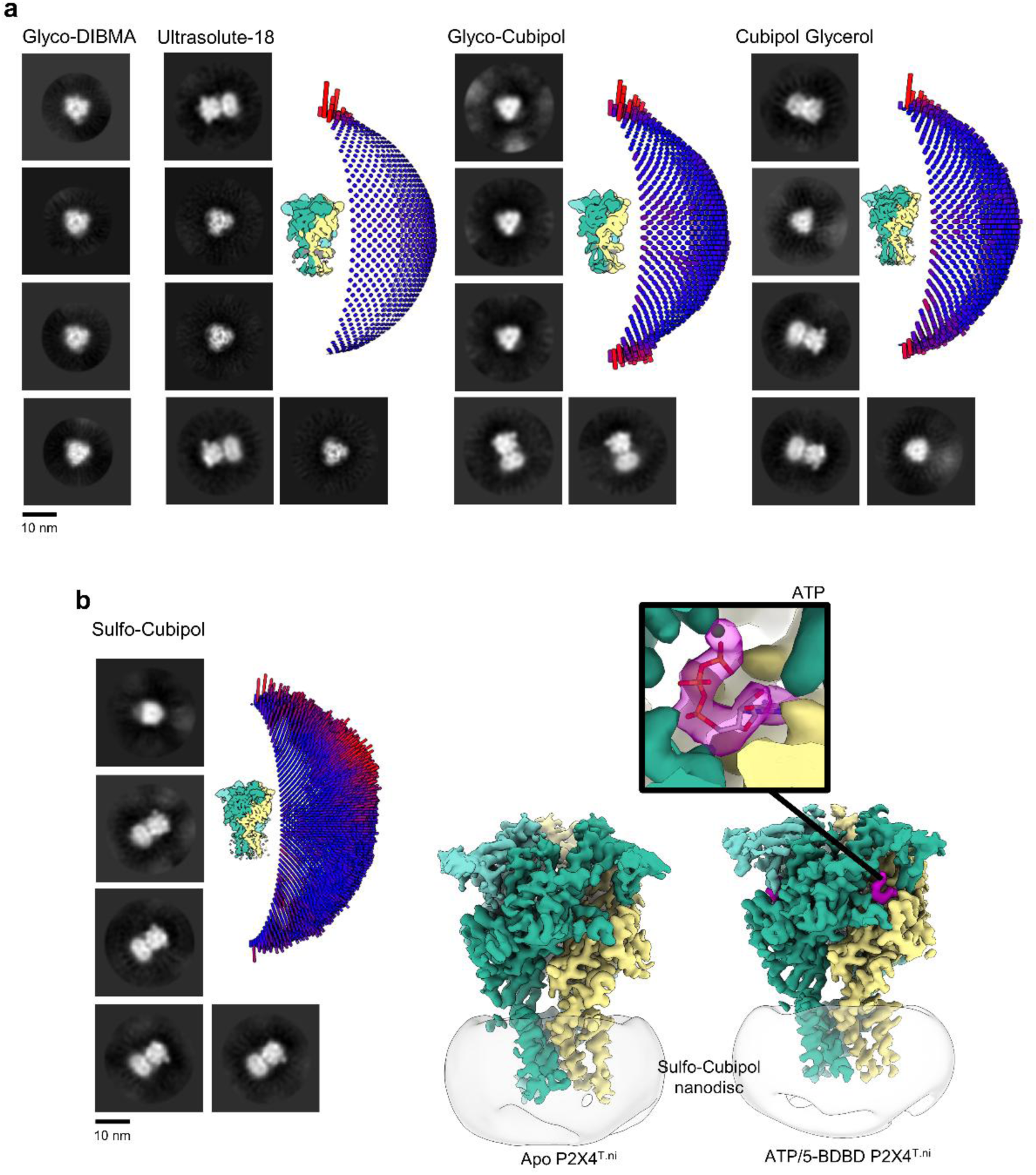
Cryo-EM screening of P2X4^T.ni^ in different polymers. **a**. Representative 2D classes obtained from the data sets of P2X4^T.ni^ in Glyco-DIBMA, Ultrasolute-18, Glyco-Cubipol, and Cubipol Glycerol. Glyco-DIBMA shows almost absolute preferential orientation. For all other polymers, a reconstruction together with an angular distribution histogram is shown next to the 2D classes. **b**. P2X4^T.ni^ in Sulfo-Cubipol was chosen for a large Cryo-EM data set. As above, 2D classes and angular distribution plots are shown on the left. High-resolution cryo-EM reconstructions of Apo P2X4^T.ni^ or in complex with ATP/5-BDBD are shown on the right. Density corresponding for ATP is highlighted in magenta, and shown in more detail in the close-up panel.

Among the tested conditions, Cubipol Glycerol and Sulfo-Cubipol produced the best-performing samples, yielding a higher fraction of side views (Fig. 7a). We attribute this to the reduced overall charge of the polymer nanodiscs, as part of the carboxylic groups of the polymer’s backbone are neutralized with either the sulfo-betaine or glycerol side chains. Interestingly, DDM/CHS solubilized P2X4 results in an orientation distribution more biased towards the particle’s side view, further stressing that the choice of solubilization agent can influence the orientation of the Cryo-EM samples ^38^. We chose to move forward with Sulfo-Cubipol, which resulted in a reconstruction of an overall resolution of 3.5 Å resolution (Fig. 7b, Extended Data Fig. 11a, Table S2). We anticipate that P2X4^Tni^ solubilized in Glycerol-Cubipol would result in a reconstruction of similar quality. All transmembrane helices are resolved, with the channel in the expected closed state. To further test the Cryo-EM compatibility of Sulfo-Cubipol solubilized P2X4^Tni^, we collected an additional data set where the protein was pre-incubated with P2X4’s agonist ATP, as well as the antagonist 5-BDBD. The complex could be resolved to an overall resolution of 2.9 Å (Fig. 7b, Extended Data Fig. 11a, Table S2). Clear additional density corresponding to MgATP is present at the agonist-binding site (Fig. 7b). As in the apo structure, the transmembrane helices are resolved, showing the channel again in its closed state. This is consistent with the presence of the additional antagonist 5-BDBD.

Unfortunately, we could not unequivocally identify the molecule’s binding site. We hypothesize that this is due either to 5-BDBD binding to the worst resolved areas of the protein (e.g. the cytoplasmic end of the nanodisc) or due to non-stoichiometric binding, particularly if the ligand binds somewhere along the symmetry axis of the protein. Together, these results demonstrate that although P2X4^Tni^ represents a biochemically suboptimal construct, upscaling with the right copolymers allows it to be structurally resolved to high resolution.

## Discussion

In this work, we introduced a semi-automated, high-throughput system for screening copolymers in membrane protein solubilization. The platform uses 96-well plates preloaded with lyophilized, pre-buffered copolymers, to which clarified lysate is directly added, and a complementary magnetic bead plates with dehydrated, storable beads for rapid affinity purification. This design enables parallel testing with minimal preparation, reducing time, labor, and variability compared to manual protocols. Applying the method to 14 human membrane proteins, we consistently obtained pure proteins embedded in native nanodiscs. Yields with next-generation copolymers rival state-of-the-art detergents, underscoring the potential of this technology to transform MP biochemistry.

Miniaturization lowers consumption of cells, beads, and copolymers, while faster processing reduces protein degradation, especially important for fragile human MPs. A single 96-well plate can accommodate three full NativeMP^TM^ copolymer sets, enabling screening of up to three targets in parallel. With a runtime of 45 min per plate (starting from lysate), ∼50 targets can be screened per week, or ∼2,500 per year on a single robot, with genome-scale coverage achievable using parallel instruments.

Although implemented in 96-well plates, the format is flexible: 384-well plates could further boost throughput, while 24-well plates would yield larger protein quantities for downstream assays. Ultimately, the choice of copolymer for upscaling depends not only on yield but also on compatibility with the intended application, for example, avoiding styrene-containing copolymers in spectroscopy, or using sulfo-betaine copolymers for complexes requiring divalent cations.

Independent of the specific copolymer finally chosen to purify the target MPs, several key biochemical characterization assays are compatible, and even only possible, using copolymer-based native nanodiscs. For instance, we and others have shown that the copolymer technology is fully compatible with mass spectrometry, either applied to protein work ^16,26^ or to lipidomics ^15^. Mass photometry has also been used to characterize the oligomerization state MPs in their native environment ^40^. On the therapeutics-development side, Dodge and colleagues ^41^ recently demonstrated that copolymer-solubilized MPs can be used in yeast display assays to raise specific nanobodies against MPs. Through our collaborations, we have also seen excellent preliminary results with affinity-selection mass spectrometry as well as DNA-encoded libraries.

To further facilitate functional studies of native membrane proteins, we developed biotinylated versions of many NativeMP™ copolymers. By directly modifying the polymer belt rather than the protein itself, this approach enables the immobilization of fully native MPs without genetic or chemical tags. We demonstrate this using P2X4^HEK^, which was captured onto a streptavidin chip for grating-coupled interferometry analysis. A key advantage is that no additional antibodies are required for immobilization, simplifying the assay and increasing sensitivity. Importantly, the same biotinylated nanodiscs can also be used in solution-based assays, providing flexibility across platforms such as fluorescence spectral shift, surface plasmon resonance (SPR), or biolayer interferometry (BLI).

Although Kd values obtained from spectral shift and GCI differed (15.6 µM vs. 1.3 µM for BAY-1797; 5.4 µM vs. ∼10 µM for 5-BDBD), both assays consistently placed antagonist binding to P2X4 in the low- to sub-micromolar range. Such differences are expected given the distinct principles of solution- versus surface-based methods, which probe protein-ligand interactions under different physical constraints. Importantly, both estimates are in line with published cellular potencies, recognizing that EC₅₀ values reflect downstream signaling amplification rather than equilibrium binding. The convergence of results across orthogonal methods demonstrates that copolymer-stabilized nanodiscs preserve the structural integrity and pharmacological competence of P2X4.

A special application is the structural determination of MPs, which has been significantly delayed behind soluble proteins. The majority of the structures deposited to the PDB come from detergent-solubilized samples, with a much smaller fraction coming from proteins either reconstituted into MSP-based or other protein-scaffolded nanodiscs ^3^ (e.g. Salipro particles). However, other than co-translationally inserted proteins ^42,43^, this also requires the use of detergents. The detergent dominance here is not unexpected, as they have been the standard for membrane protein work for the last decades. Nevertheless, examples exist of structures in native nanodiscs formed with most copolymer families. However, systematic analyses of different copolymers for Cryo-EM are rarely reported, except for specific examples ^44^. Here, we demonstrated that several copolymers could lead to a high-resolution structure of P2X4, with our Sulfo-Cubipol structures reaching 2.9 Å resolution. Chemically modifying the copolymers had a strong influence on the orientation bias of the protein, suggesting that the copolymer’s identity can be used to fine-tune the protein’s orientation. In turn, this also demonstrates that a specific Cryo-EM copolymer screening is likely required for each target. We demonstrated here that, in principle, no additional upscale purification is required for this, as elutions from the magnetic bead plate could be directly taken into the microscope. One could envision further automation of this, where the plate elutions are directly transferred to a state-of-the-art vitrification instrument such as Chameleon (SPT Labtech), Vitrojet (CryoSol) or CryoWriter^45^. The latter allows to sample directly from 96-well plates, allowing for a fully automated protocol.

## Acknowledgements

We thank Dr. Thomas Heidler, Dr Saba Shahzad, and Sandeep Signh for their assistance during Cryo-EM data collection at Forschungszentrum Jülich.

## Data Availability

All maps are deposited to EMDB and have the the following accession numbers: P2X4^HEK^ in Cubipol Glycerol, EMD-59335; P2X4^T.ni^ in Ultrasolute-18, EMD-59364; P2X4^T.ni^ in Glyco-Cubipol, EMD-59363; P2X4^T.ni^ in Cubipol Glycerol, EMD-59361; P2X4^T.ni^ in Sulfo-Cubipol, EMD-59333; and P2X4^T.ni^/ATP in Sulfo-Cubipol, EMD-59354. Atomic models for P2X4^T.ni^ in Sulfo-Cubipol and P2X4^T.ni^/ATP in Sulfo-Cubipol have the PDB codes pdb_000033BX and pdb_000033CQ respectively.

## Methods

### Construct design

Human full-length SLC1A6 (P48664), SLC2A1 (P11166), SLC15A4 (Q8N697), CLDN1 (O95832), CLDN3 (O15551), CLDN4 (O14493), CD68 (P34810), CD163 (Q86VB7), EGFR (P00533), P2X4 (Q99571), and FAS (P25445) were synthesized with a C-terminal Rho1D4 tag connected via a GSSG linker, codon-optimized for HEK293 expression (GeneArt, Thermo Fisher), and cloned into pcDNA3.4-TOPO (Thermo Fisher). HTR7 (P34969) and ADRB2 (P07550) followed the same design but carried N-terminal HA-Flag or HA-Flag-HRV3C tag, respectively, and were codon-optimized for *T.ni* cells and cloned into pOET2 vector (Oxford Expression Technologies, OET) like the AMY3R complex comprised CalcR (P30988; N-terminal HA-Flag-HRV3C, C-terminal GSSG-Rho1D4) and Ramp3 (O60896; N-terminal HA-His), Truncated hERG (Q12809; Δ141-350, 871-1005) carried a C-terminal GSSG-Prescission-Rho1D4 tag and was inserted into pcDNA3.4-TOPO.

Human full length P2X4 (NP_002551.2; residues 1-388, mutations R75N, R184N, T307E) was codon-optimized for expression in insect cells (GeneArt, Thermo Fisher) and subcloned (EcoRI/BamHI) into pOET2 expression vector with a C-terminal GSSG-Rho1D4 tag.

### Expression of GPCRs in *T.ni* insect cells

GPCRs were expressed using the flashBAC™ technology (OET). Sf9 cells (Sf-900™, Gibco) were co-transfected with flashBAC ULTRA™ DNA and pOET2 carrying the gene of interest using baculoFECTIN II (OET) in serum-free Insect-XPRESS™ medium (Lonza). After 5 days at 27 °C, the supernatant containing recombinant virus (P0) was collected and stored in the dark at 4 °C. For amplification, Sf9 suspension cultures (2 x 10⁶ cells/mL) were infected with P0 or P1 virus and incubated 4-5 days at 27 °C, 110 rpm. Clarified supernatants were stored at 4 °C. For protein expression, *T.ni* cells (High Five™, Gibco) (2.5 x 10⁶ cells/mL) were infected with P1 or P2 virus and cultured 48 h at 27 °C, 110 rpm. Pellets were submerged in 1mL buffer (20 mM HEPES, 10 mM NaCl, 2 mM EDTA containing protease inhibitors) and stored at −80 °C.

### Transient expression in Expi293F (HEK) cells

Expi293F cells (Gibco) were transiently transfected with plasmid DNA using 40 kDa linear polyethylenimine (PEI; PolySciences). Cells were cultured at 2.5 x 10⁶ cells mL in serum-free SMM 293-TII medium (Sino Biological) supplemented with 0.1% Pluronic F-68. DNA/PEI complexes (1.25 µg/mL DNA; 6.2 µg/mL PEI) were pre-incubated for 20 min at RT before addition to 300 mL cultures. Transfected cells were maintained at 37 °C, 6% CO₂, and 120 rpm. Sodium butyrate (10 mM) was added 20 h post-transfection, and cells were harvested after 72 h by centrifugation (2,000 rpm, 15 min, 4 °C). Pellets were submerged in 1mL buffer (20 mM HEPES, 10 mM NaCl, 2 mM EDTA containing protease inhibitors) and stored at −80 °C.

### Manual screening (6.5h)

Cell pellets (HEK or *T.ni*) expressing the target protein were thawed in lysis buffer (1:10, v/v) at RT and sonicated on ice (three cycles, ∼5 min each). Lysates were clarified by centrifugation (9,000 x g, 30 min, RT). Supernatants (1 mL aliquots) were ultracentrifuged (60,000 x g, 45 min, RT), and membrane pellets were weighed and resuspended in 1 mL of 2.5% polymer solution. Solubilization was carried out for 2 h at RT. After re-centrifugation (60,000 x g, 45 min, RT), insoluble fractions were weighed to determine solubilization efficiency. Supernatants were diluted (1:10, v/v) in protein buffer and incubated overnight with 50 µL affinity magnetic beads (MagBeads) at RT. Beads were washed three times with protein buffer, and bound proteins were eluted using 2 mg/mL Rho peptide, 1 mg/mL FLAG peptide, or 50 mM biotin (Rho1D4, FLAG, or StrepTactinXT resin). Eluates were analyzed by DLS, nanoDSF, SDS-PAGE, and Western blotting.

### Classical versus semi-automated copolymer workflow

The classical workflow (Fig. 1a) for membrane protein extraction involves cell lysis, clarification by centrifugation, membrane pelleting, copolymer-mediated solubilization at 4 °C for up to 16 h, reclarification, and binding to affinity resin and purification, which requires ∼1200 min.

In the semi-automated workflow (Fig. 1b,c), clarified lysates (9000 x g supernatant, 9kS) are applied directly to lyophilized copolymer plates (NativeMP^TM^, 25 mg/well), bypassing membrane pelleting and ultracentrifugation. Rho1D4 magnetic beads (30 µL; PureHT^TM^ plate) are stored dehydrated and rehydrated during lysate clarification, minimizing preparation time. All steps are executed on a KingFisher Flex system using fast-mixing mode with a single tip comb for bead transfer and agitation. After loading, solubilization is completed within 5 min at room temperature (RT) or up to 37 °C, replacing overnight incubation. Binding to Rho1D4 beads occurs simultaneously. Beads are washed in three plates containing 1 mL wash buffer (20 mM HEPES, 150 mM NaCl, pH 7-8) for 5 min each and eluted in 50 µL elution buffer (20 mM HEPES, 150 mM NaCl, 2 mg/mL Rho peptide, pH 7-8).

Eluates were directly applied to downstream assays, including DLS, nDSF, SDS-PAGE, Western blotting, and Cryo-EM. A brief post-elution centrifugation (2 min at 500 x g, then 12,000-50,000 x g for 10 min) removes residual aggregates or beads.

For typical experiments, 7-10 mL of pelleted cell suspension (2.5-4 x 10⁶ cells/mL) were lysed in protein buffer and clarified for 45 min at 9000 x g. One mL supernatant per well was used; screening 32 copolymers required ∼250–300 mL culture adjusted to 32 mL total supernatant. The semi-automated format is fully scalable and allows parallel screening of multiple copolymers or conditions in a single run.

### Upscale of P2X4

P2X4 cell pellets were thawed and resuspended in lysis buffer at RT (20 mM HEPES, pH 7.5, 150 mM NaCl, protease inhibitor cocktail) and subjected to sonication (3x 5 min). Lysates were clarified by centrifugation (9,000 x g for 45 min at RT), supernatant was incubated for 30 min with apyrase (1:100 dilution in divalent cation-containing buffer) to remove protein-bound ATP. Membranes were subsequently isolated by ultracentrifugation (100,000 x g for 45 min), collected, weighed, and solubilized at RT using 2.5% of copolymer in solution dissolved in 15 mL protein buffer per gram of membrane. The suspension was sonicated (3x 3 min) to enhance solubilization, followed by incubation for ∼2 h. Insoluble membranes were removed by ultracentrifugation at 100,000 × g for 45 min. Copolymer stabilized P2X4 was captured by incubation with Rho MagBeads for 2 h at RT, washed extensively using protein buffer, and eluted using 2 mg/mL Rho peptide in protein buffer. Eluates were concentrated using a 50-kDa molecular weight cutoff Amicon concentrator and subjected to size-exclusion chromatography (SEC) on a Superose 6 Increase 10/300 GL column. Peak fractions were collected, concentrated, and used for subsequent biophysical and biochemical characterization.

### Protein quality control

#### SDS-PAGE analysis and membrane protein quantification

MPs were analyzed by SDS-PAGE and Coomassie staining. Band intensities were quantified in ImageJ and integrated after background subtraction. Protein fluorescence at 330 nm was measured on a NanoTemper Panta Device. All values were normalized to the strongest signal (100%) and grouped by copolymer family and visualized as box plots to represent the distribution of protein yields for each target.

#### DLS and NanoDSF

Copolymer-stabilized MPs were spinned at high speed (50,000 x g, 10 min) before subjecting to the DLS and nanoDSF measurement on a NanoTemper Prometheus Panta using 10 µL capillaries. Thermal unfolding was monitored from 25-95 °C, at 3 °C/min with DLS enabled; F350/F330 ratios and first derivatives yielded T_onset_ and T_m_, while DLS tracked temperature-dependent size and aggregation. Isothermal DLS holds (25-95 °C) assessed size stability (Z-average, PDI). Data were analyzed in Prism and NanoTemper software.

#### Biochemical characterization using fluorescence shift assay

Binding studies were performed using fluorescence shift measurements on a Monolith X instrument (NanoTemper Technologies). Copolymer-stabilized P2X4 was fluorescently labeled according to the manufacturers protocol using the NT-L021 Protein Labeling Kit (Lysine-reactive; NanoTemper Technologies). For the Monolith X measurement 40 nM labelled protein was mixed with each ligand dilution (16 of 1:1 dilutions), thus resulting in a final protein concentration of 20 nM. After 30 min incubation, the samples were loaded into Monolith X by using the Monolith glass Capillaries (cat# MO-K022, NanoTemper Technologies).

BAY 1797 was serial diluted in MST buffer (20 mM HEPES, pH 7.5, 150 mM NaCl, 0.005% Tween-20) and mixed with P2X4 before loading into the capillaries (NanoTemper Technologies). Fluorescence shifts (ΔRatio 670/650 nm) were recorded at 25 °C using 40% LED excitation and medium laser power. Thermophoretic movement was monitored for 30 s following IR-laser activation. Data were analyzed with MO. Affinity Analysis software (NanoTemper Technologies). Dissociation constants (K_d_) or IC_50_ were determined by fitting with a nonlinear regression model. Each experiment was performed in triplicates. Equivalent binding assays were also conducted for the ligand 5-BDBD under identical conditions.

#### Biochemical characterization using grating-coupled interferometry

Binding kinetics of the P2X4 receptor were characterized using grating-coupled interferometry (GCI; WAVEsystem, Creoptix). Full-length P2X4 reconstituted in Ultrasolute^TM^ Amphipol 18 (U18) biotin nanodiscs was immobilized on streptavidin capture chips (PCH-STA), while control copolymer-host lipid nanodisc with U18 biotin served as negative controls to account for nonspecific interactions. WaveRAPID analysis with 5-BDBD were performed at a final concentration 20 µM, injections were carried at 50 µl/min with an association phase of 25 s and dissociation phase of 400 s in 20 mM HEPES, 150 mM NaCl, 1% DMSO, 0.05% TWEEN, pH 7.5. Binding analysis was fitted using a conformational-change model. Multicycle analyses were performed with the analyte BAY-1797, therefore injections in a twofold dilution series across a 39 nM-20 µM concentration range at a flow rate of 12.5 µl min with 60 s association and 60 s dissociation phase in 20 mM HEPES, 150 mM NaCl, 1% DMSO, pH 7.5 were conducted to the system. Data were corrected against the control copolymer-host lipid nanodisc reference, a DMSO calibration, buffer blanks and globally fitted using a 1:1 binding model and conformation aI-change model. The resulting rate constants (k_on_, k_off_) and derived dissociation constants (K_d_) were derived from the global fits and reported as mean values from duplicate measurements.

#### Cryo-EM sample preparation, data collection and analysis

Copolymer solubilized P2X4 samples were concentrated to 1.3 - 2.0 mg/mL before grid vitrification. The King Fisher purified samples had lower concentrations due to the overall small size of the purification. For samples with ligands, a preincubation with 100 µM of the desired small molecule before freezing was performed. For all samples, a 3.5 µL drop was applied onto a freshly glow-discharged Cryo-EM grid: UltraAU Foil 1.2/1.3 for manually purified P2X4 or Quantifoil R1.2/1.3 for the King Fisher purifications. The sample chamber was kept at 15 °C and 95% humidity. The excess sample was blotted (force −5, 3.5 s) and the grid was vitrified in a Vitrbot IV instrument using liquid ethane.

The samples were then transferred into a cryo microscope for further analysis. In total, we used two different instruments for this purpose. (i) A Talos Arctica equipped with an XFEG operated at 200 kV, a K3 detector and a BioQuantum energy filter, or (ii) a Titan Krios G4i equipped with an XFEG operated at 300 kV, a K3 detector coupled to a BioContinuum energy filter or a Falcon 4 detector. In all cases, the energy filter was set to a slit width of 20 eV, and the K3 detector was used in CDS super-resolution mode. Table S1 summarizes the microscope and detector combination, exposure settings, and data set sizes for each sample

While the number of images differs for every dataset, we aimed at 1,000 – 1,500 images to determine whether a sample could lead to a successful 3D reconstruction. All micrographs were collected at defocus values between −1.0 and −3.0 µm, with a physical pixel size of ∼ 0.8 Å/px (see Table S1). All single particle work was done with Relion ^46^. Movies were aligned and dose-corrected using either Relion’s MotionCor implementation ^47^ or MotionCor3 (https://github.com/czimaginginstitute/MotionCor3). CTF estimation was performed with CTFFIND 4 ^48^. Particles were automatically selected using the generalized model of Topaz ^49^. The particles were extracted into 384px boxes, and downsampled to 96px. After several rounds of 2D classification, good particles were selected to re-train a topaz model, which was subsequently used to pick particles again. The new particles were extracted in 384px boxes, and downsampled as required. After an initial cleaning with 2D classification, we determined whether enough orientations were visible to allow for 3D work. An initial 3D model was generated using stochastic gradient descent ^50^, and several rounds of 3D classification followed by 3D refinement were used to obtain the final structure. Map regularization was done using Blush ^51^. For P2X4^T.ni^ Sulfo-Cubipol and P2X4^HEK^, an additional round of Bayesian polishing ^47^ followed by CTF refinement ^52^ was done in order to improve the resolution of the reconstructions. We used cryoSieve ^53^ as a final cleaning step for these samples, resulting in reconstructions of 3.5 Å for Apo P2X4^T.ni^ and 2.9 Å for P2X4^T.ni^ bound to ATP/5-BDBD. Additionally, we built atomic models for these two structures. In both cases, we used ModelAngelo^54^ to create to starting model. The models were completed in Coot^55^ and further refined using Rosetta’s iterative local rebuilding^56^. The final models were refined using Servalcat^57^.

#### Polymer overview

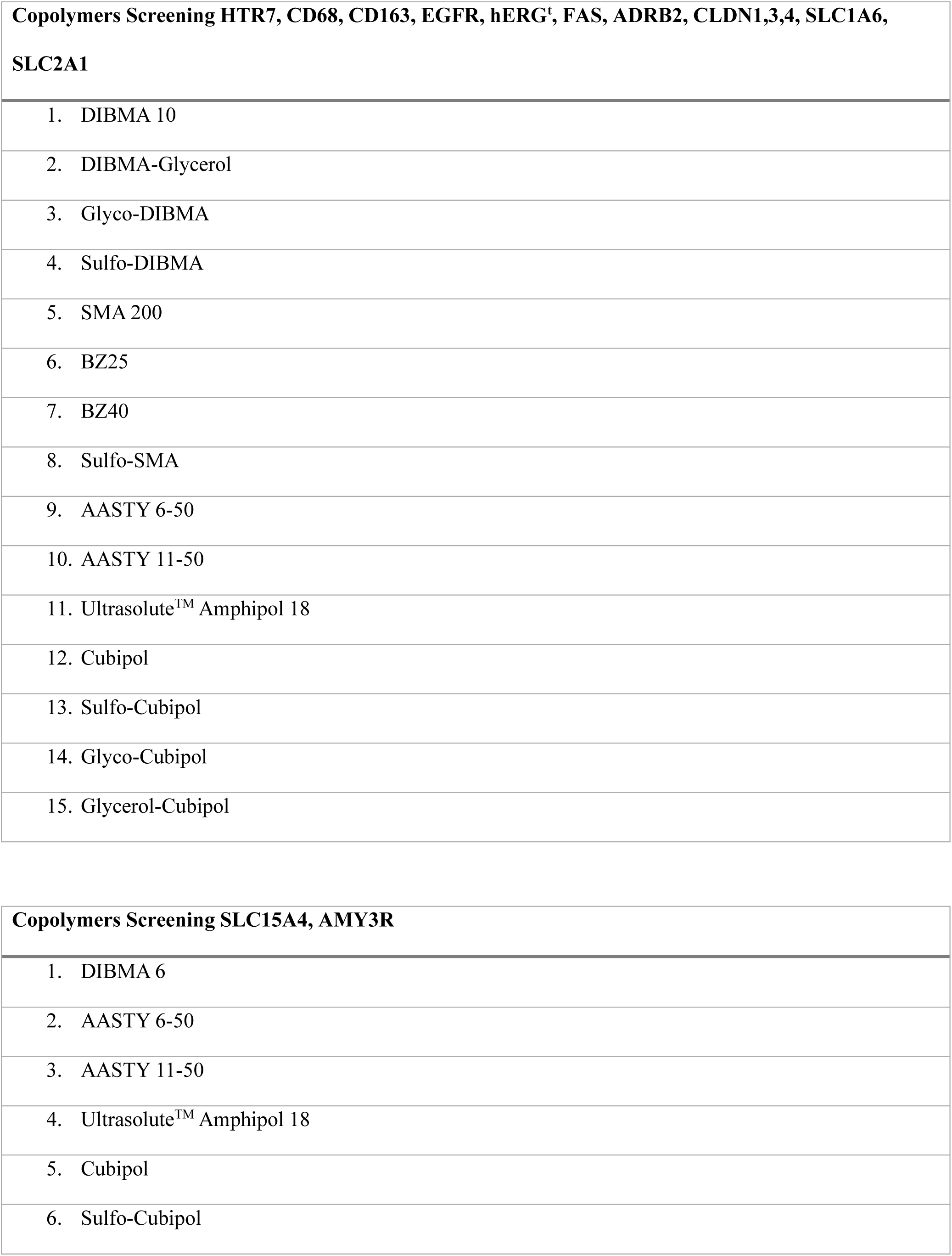
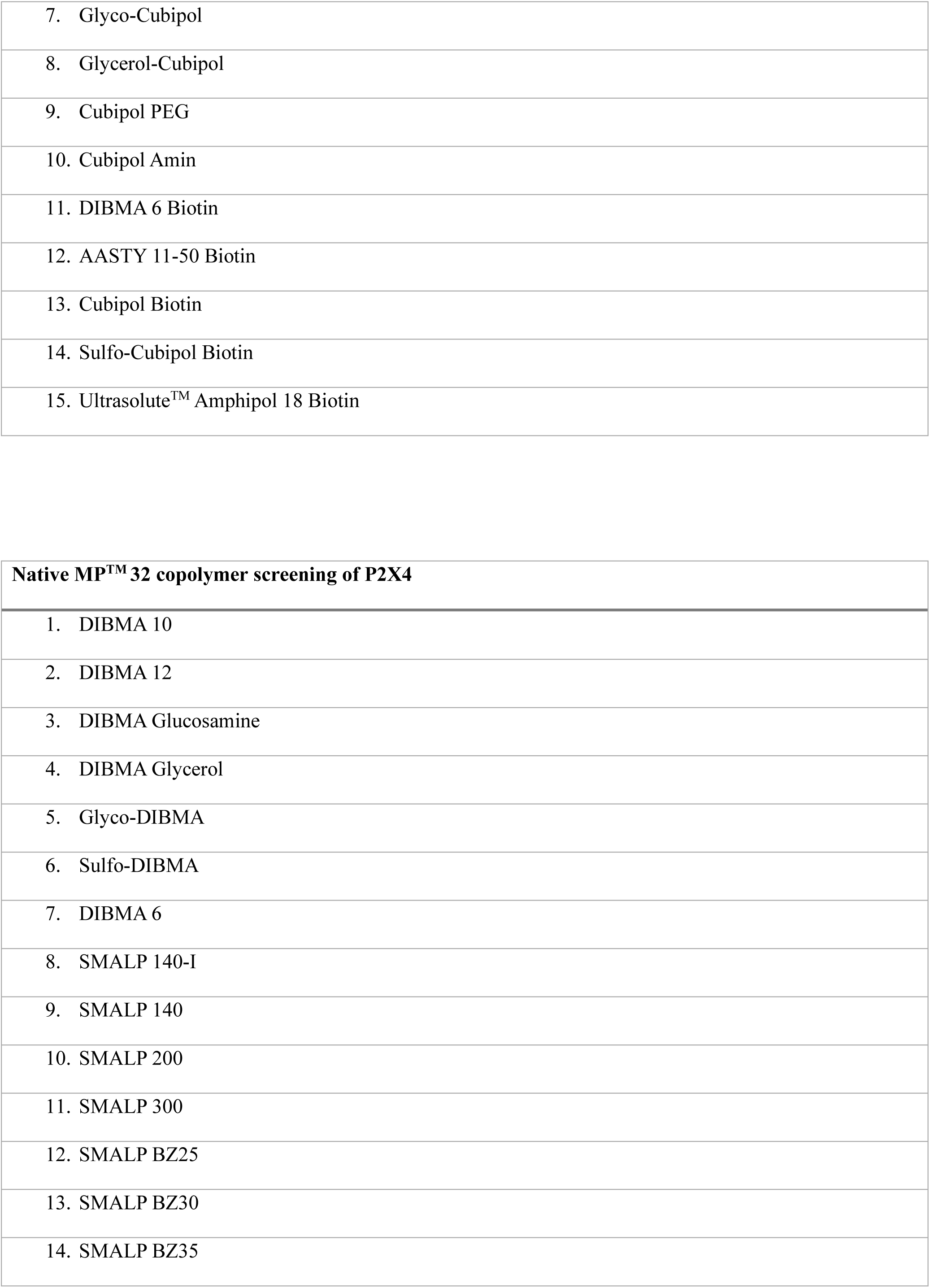
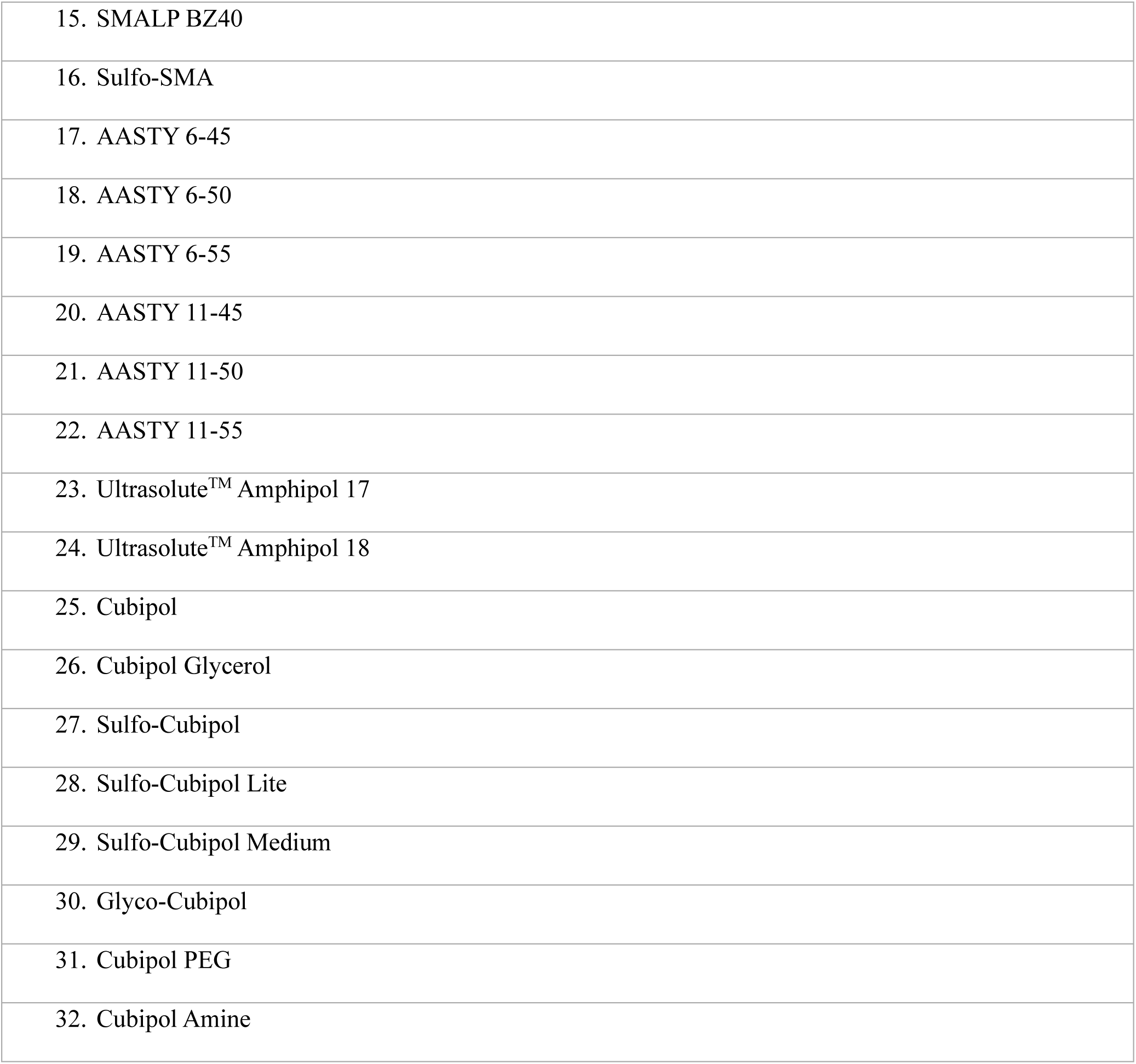

| **Copolymers Screening HTR7, CD68, CD163, EGFR, hERG<sup>t</sup>, FAS, ADRB2, CLDN1,3,4, SLC1A6, SLC2A1** |  |
| --- | --- |
| 1. | DIBMA 10 |
| 2. | DIBMA-Glycerol |
| 3. | Glyco-DIBMA |
| 4. | Sulfo-DIBMA |
| 5. | SMA 200 |
| 6. | BZ25 |
| 7. | BZ40 |
| 8. | Sulfo-SMA |
| 9. | AASTY 6-50 |
| 10. | AASTY 11-50 |
| 11. | Ultrasolute™ Amphipol 18 |
| 12. | Cubipol |
| 13. | Sulfo-Cubipol |
| 14. | Glyco-Cubipol |
| 15. | Glycerol-Cubipol |

| **Copolymers Screening SLC15A4, AMY3R** |  |
| --- | --- |
| 1. | DIBMA 6 |
| 2. | AASTY 6-50 |
| 3. | AASTY 11-50 |
| 4. | Ultrasolute™ Amphipol 18 |
| 5. | Cubipol |
| 6. | Sulfo-Cubipol |

|  |
| --- |
| 7. Glyco-Cubipol |
| 8. Glycerol-Cubipol |
| 9. Cubipol PEG |
| 10. Cubipol Amin |
| 11. DIBMA 6 Biotin |
| 12. AASTY 11-50 Biotin |
| 13. Cubipol Biotin |
| 14. Sulfo-Cubipol Biotin |
| 15. Ultrasolute™ Amphipol 18 Biotin |

| **Native MP™ 32 copolymer screening of P2X4** |
| --- |
| 1. DIBMA 10 |
| 2. DIBMA 12 |
| 3. DIBMA Glucosamine |
| 4. DIBMA Glycerol |
| 5. Glyco-DIBMA |
| 6. Sulfo-DIBMA |
| 7. DIBMA 6 |
| 8. SMALP 140-I |
| 9. SMALP 140 |
| 10. SMALP 200 |
| 11. SMALP 300 |
| 12. SMALP BZ25 |
| 13. SMALP BZ30 |
| 14. SMALP BZ35 |
| 15. SMALP BZ40 |
| 16. Sulfo-SMA |
| 17. AASTY 6-45 |
| 18. AASTY 6-50 |
| 19. AASTY 6-55 |
| 20. AASTY 11-45 |
| 21. AASTY 11-50 |
| 22. AASTY 11-55 |
| 23. Ultrasolute™ Amphipol 17 |
| 24. Ultrasolute™ Amphipol 18 |
| 25. Cubipol |
| 26. Cubipol Glycerol |
| 27. Sulfo-Cubipol |
| 28. Sulfo-Cubipol Lite |
| 29. Sulfo-Cubipol Medium |
| 30. Glyco-Cubipol |
| 31. Cubipol PEG |
| 32. Cubipol Amine |

