## Supplementary Material for "Membrane Proteins at Scale: Automated Copolymer Nanodisc Purification for Structure and Function"

### Extended Data

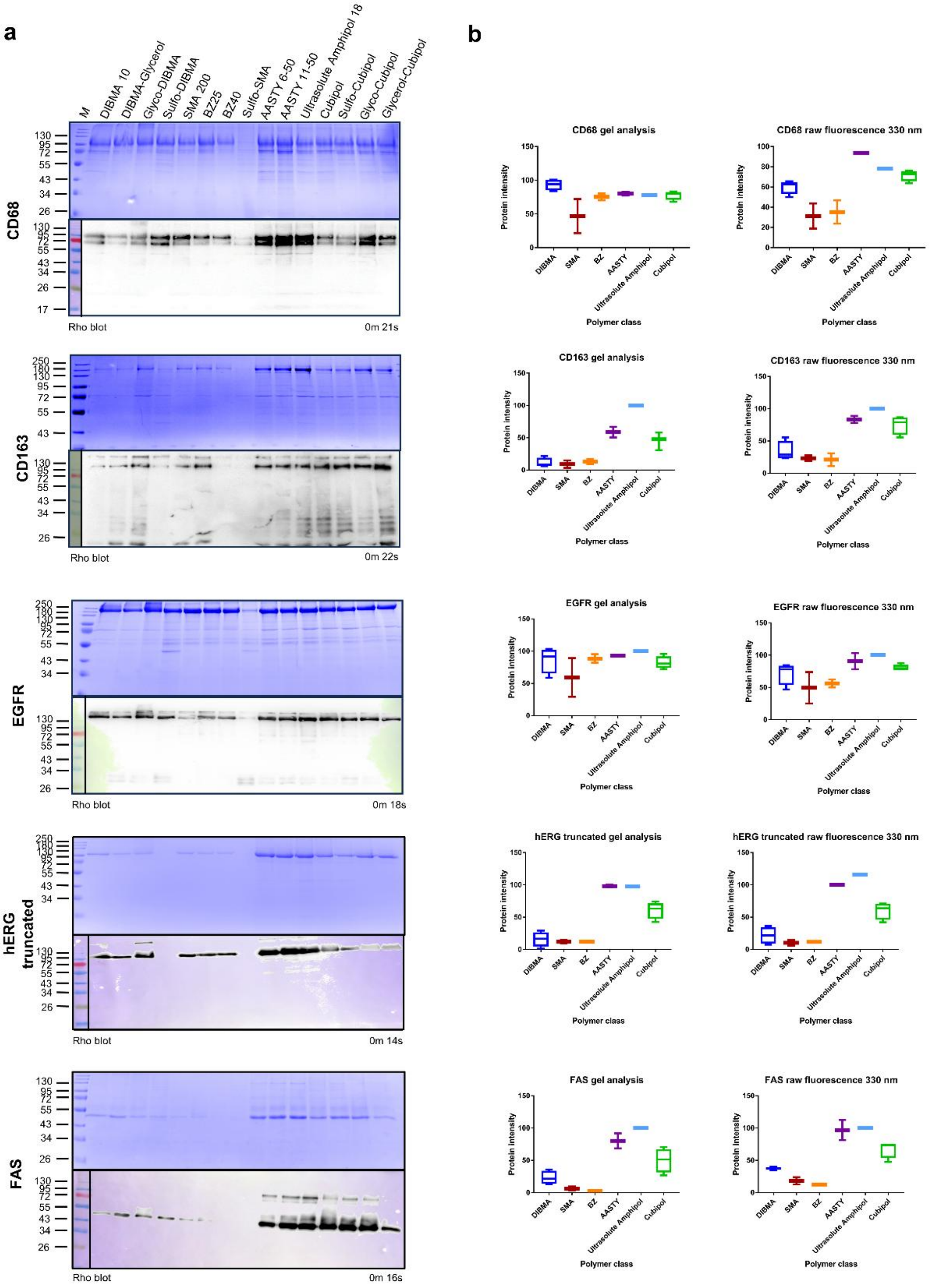


**Extended Data Figure 1. Copolymer screening of CD68, CD163, EGFR, hERG$^t$, and FAS**

a. Coomassie-stained SDS–PAGE and corresponding Western blots of CD68, CD163, EGFR, hERG$^t$, and FAS solubilized and purified in nanodiscs formed with 15 different copolymers. For Figure 2, only the best-performing copolymer eluates were shown, while here the full screening dataset is displayed. b. Quantitative analysis of the same screens by densitometric gel analysis and intrinsic tryptophan fluorescence at 330 nm. Values were normalized to the strongest signal within each target dataset (set to 100%) and plotted to illustrate recovery and solubilization efficiency across copolymer families.

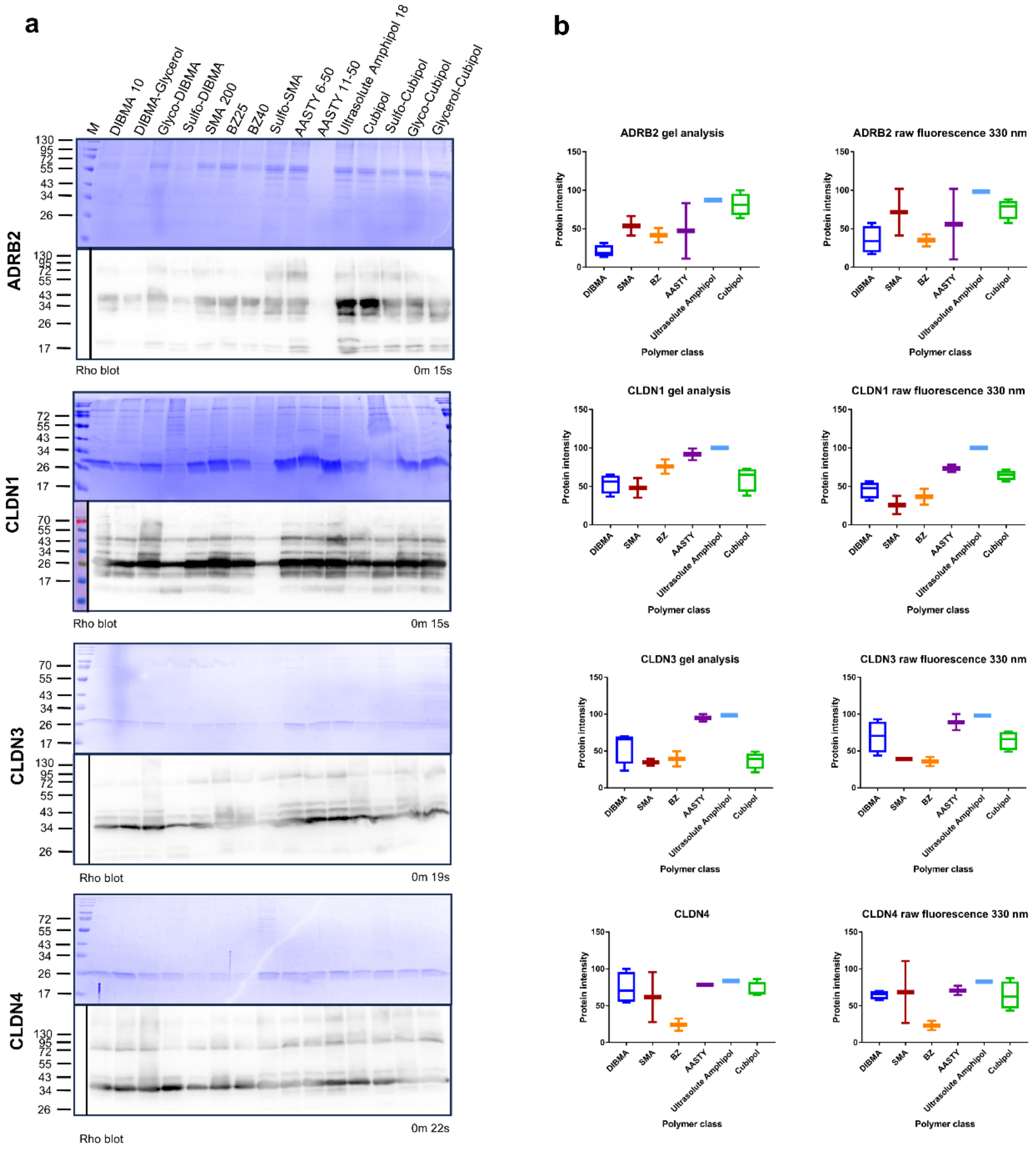


**Extended Data Figure 2. Copolymer screening of ADRB2, CLDN1, CLDN3, and CLDN4**

a. Coomassie-stained SDS–PAGE and corresponding Western blots of ADRB2, CLDN1, CLDN3, and CLDN4 solubilized and purified in nanodiscs formed with 15 different copolymers. For Figure 2, only the best-performing copolymer eluates were shown, while here the full screening dataset is displayed. b. Quantitative analysis of the same screens by densitometric gel analysis and intrinsic tryptophan fluorescence at 330 nm. Values were normalized to the strongest signal within each target dataset (set to 100%) and plotted to illustrate recovery and solubilization efficiency across copolymer families.

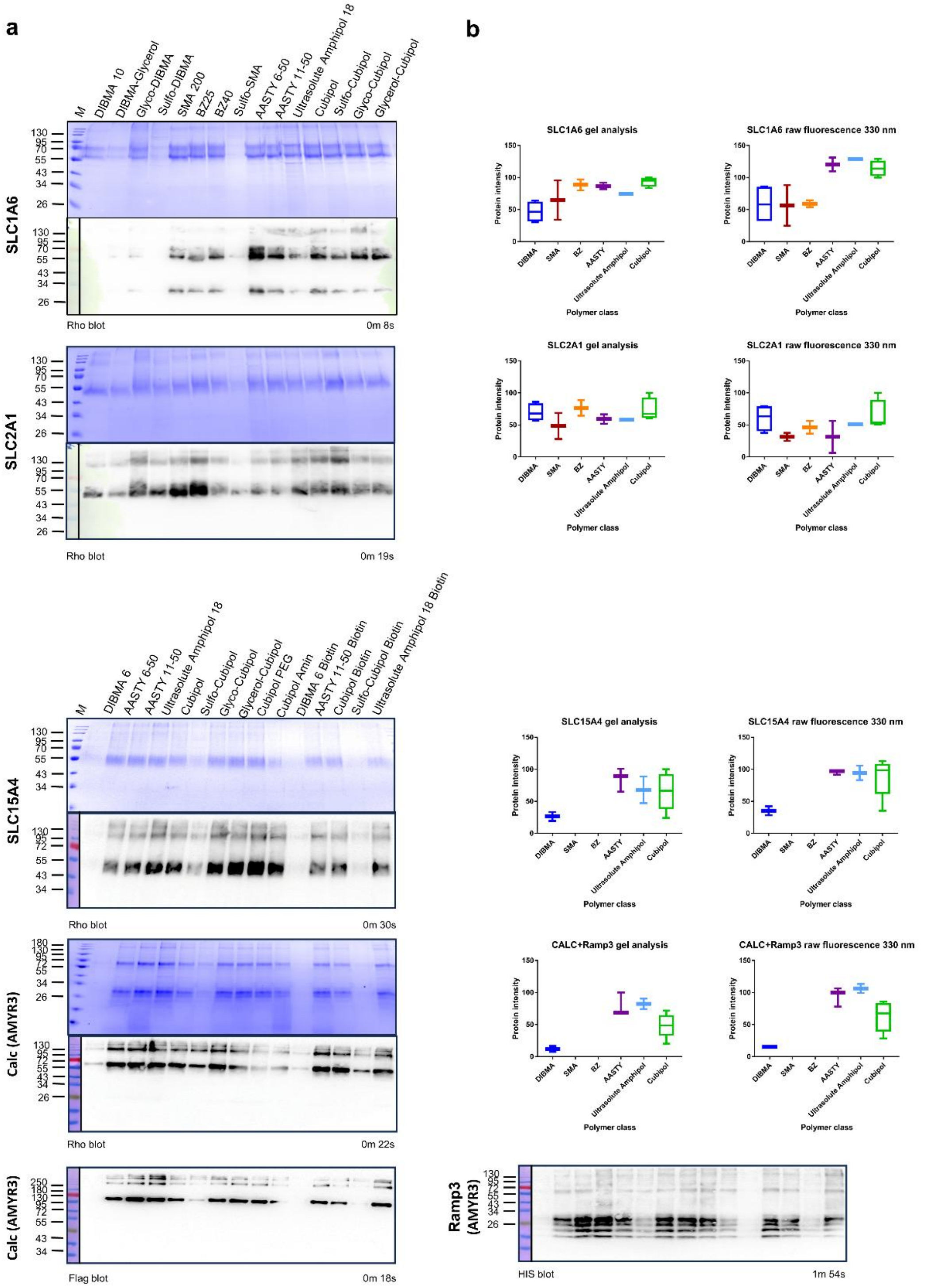


**Extended Data Figure 3. Copolymer screening of SLC1A6, SLC2A1, SLC15A4, AMYR3 (Calc+Ramp3)**

a. Coomassie-stained SDS–PAGE and corresponding Western blots of SLC1A6, SLC2A1, SLC15A4, and AMYR3 (co-expressed with Ramp3) solubilized and purified in nanodiscs formed with 15 different copolymers. For Figure 2, only the best-performing copolymer eluates were shown, while here the full screening dataset is displayed. b. Quantitative analysis of the same screens by densitometric gel analysis and intrinsic tryptophan fluorescence at 330 nm. Values were normalized to the strongest signal within each target dataset (set to 100%) and plotted to illustrate recovery and solubilization efficiency across copolymer families.

**a**

| Coomassie Gel analysis | DIBMA | SMA | BZ | AASTY | Ultrasolute Amphipol | Cubipol |
|---|---|---|---|---|---|---|
| Number of values | 52 | 24 | 24 | 30 | 16 | 64 |
| Minimum | 0,94 | 3,33 | 1,6 | 11,19 | 47,16 | 20,03 |
| 25% Percentile | 16,47 | 14,86 | 16,15 | 66,85 | 75,7 | 47,55 |
| Median | 35,95 | 29,72 | 41,21 | 82,37 | 89,73 | 64,68 |
| 75% Percentile | 68,57 | 65,12 | 80,27 | 93,04 | 100 | 80,39 |
| Maximum | 103,7 | 95,67 | 97,08 | 100,6 | 100 | 100 |
| Mean | 44,84 | 38,48 | 46,8 | 77,43 | 86,14 | 63,34 |
| Std. Deviation | 31,14 | 29,13 | 32,51 | 20,8 | 16,16 | 21,57 |
| Std. Error of Mean | 4,319 | 5,947 | 6,636 | 3,797 | 4,041 | 2,696 |
| Lower 95% CI of mean | 36,17 | 26,18 | 33,07 | 69,67 | 77,53 | 57,95 |
| Upper 95% CI of mean | 53,51 | 50,78 | 60,52 | 85,2 | 94,76 | 68,72 |
| Sum | 2332 | 923,5 | 1123 | 2323 | 1378 | 4054 |

| 330nm fluorescence | DIBMA | SMA | BZ | AASTY | Ultrasolute Amphipol | Cubipol |
|---|---|---|---|---|---|---|
| Number of values | 52 | 24 | 24 | 30 | 16 | 64 |
| Minimum | 7,25 | 5,8 | 10 | 6,25 | 51,25 | 28,37 |
| 25% Percentile | 29,56 | 18,79 | 13,58 | 77,59 | 86,84 | 56,6 |
| Median | 43 | 25,8 | 29,41 | 92,65 | 100 | 72,05 |
| 75% Percentile | 62,88 | 40,83 | 46,88 | 100 | 104,2 | 85,75 |
| Maximum | 93,14 | 110,6 | 64,52 | 131,2 | 129 | 129 |
| Mean | 46,78 | 36,71 | 32,5 | 84,97 | 97,17 | 72,68 |
| Std. Deviation | 22,61 | 28,61 | 17,42 | 26,18 | 17,57 | 21,2 |
| Std. Error of Mean | 3,135 | 5,839 | 3,556 | 4,779 | 4,394 | 2,65 |
| Lower 95% CI of mean | 40,48 | 24,63 | 25,15 | 75,19 | 87,81 | 67,38 |
| Upper 95% CI of mean | 53,07 | 48,79 | 39,86 | 94,74 | 106,5 | 77,98 |
| Sum | 2432 | 881,1 | 780,1 | 2549 | 1555 | 4652 |

**b**

| Coomassie Gel analysis | | |
|---|---|---|
| **SMA vs. DIBMA** | | |
| Mann Whitney test | | |
| P value | | 0,4791 |
| Exact or approximate P value? | Exact | |
| P value summary | ns | |
| Significantly different (P < 0.05)? | No | |
| One- or two-tailed P value? | Two-tailed | |
| Sum of ranks in column A,B | 2066 , 860 | |
| Mann-Whitney U | | 560 |
| **SMA vs. BZ** | | |
| Mann Whitney test | | |
| P value | | 0,4308 |
| Exact or approximate P value? | Exact | |
| P value summary | ns | |
| Significantly different (P < 0.05)? | No | |
| One- or two-tailed P value? | Two-tailed | |
| Sum of ranks in column B,C | 549 , 627 | |
| Mann-Whitney U | | 249 |
| **SMA vs. AASTY** | | |
| Mann Whitney test | | |
| P value | <0,0001 | |
| Exact or approximate P value? | Exact | |
| P value summary | **** | |
| Significantly different (P < 0.05)? | Yes | |
| One- or two-tailed P value? | Two-tailed | |
| Sum of ranks in column B,D | 413 , 1072 | |
| Mann-Whitney U | | 113 |
| **SMA vs. Ultrasolute Amphipol** | | |
| Mann Whitney test | | |
| P value | <0,0001 | |
| Exact or approximate P value? | Exact | |
| P value summary | **** | |
| Significantly different (P < 0.05)? | Yes | |
| One- or two-tailed P value? | Two-tailed | |
| Sum of ranks in column B,E | 334 , 486 | |
| Mann-Whitney U | | 34 |
| **SMA vs. Cubipol** | | |
| Mann Whitney test | | |
| P value | | 0,0002 |
| Exact or approximate P value? | Exact | |
| P value summary | *** | |
| Significantly different (P < 0.05)? | Yes | |
| One- or two-tailed P value? | Two-tailed | |
| Sum of ranks in column B,F | 678 , 3238 | |
| Mann-Whitney U | | 378 |

| 330nm fluorescence | | |
|---|---|---|
| **SMA vs. DIBMA** | | |
| Mann Whitney test | | |
| P value | | 0,0241 |
| Exact or approximate P value? | Exact | |
| P value summary | * | |
| Significantly different (P < 0.05)? | Yes | |
| One- or two-tailed P value? | Two-tailed | |
| Sum of ranks in column A,B | 2203 , 723 | |
| Mann-Whitney U | | 423 |
| **SMA vs. BZ** | | |
| Mann Whitney test | | |
| P value | | 0,8903 |
| Exact or approximate P value? | Exact | |
| P value summary | ns | |
| Significantly different (P < 0.05)? | No | |
| One- or two-tailed P value? | Two-tailed | |
| Sum of ranks in column B,C | 581 , 595 | |
| Mann-Whitney U | | 281 |
| **SMA vs. AASTY** | | |
| Mann Whitney test | | |
| P value | <0,0001 | |
| Exact or approximate P value? | Exact | |
| P value summary | **** | |
| Significantly different (P < 0.05)? | Yes | |
| One- or two-tailed P value? | Two-tailed | |
| Sum of ranks in column B,D | 408 , 1077 | |
| Mann-Whitney U | | 108 |
| **SMA vs. Ultrasolute Amphipol** | | |
| Mann Whitney test | | |
| P value | <0,0001 | |
| Exact or approximate P value? | Exact | |
| P value summary | **** | |
| Significantly different (P < 0.05)? | Yes | |
| One- or two-tailed P value? | Two-tailed | |
| Sum of ranks in column B,E | 330 , 490 | |
| Mann-Whitney U | | 30 |
| **SMA vs. Cubipol** | | |
| Mann Whitney test | | |
| P value | <0,0001 | |
| Exact or approximate P value? | Exact | |
| P value summary | **** | |
| Significantly different (P < 0.05)? | Yes | |
| One- or two-tailed P value? | Two-tailed | |
| Sum of ranks in column B,F | 527,5 , 3389 | |
| Mann-Whitney U | | 227,5 |

**Extended Data Figure 4. Statistical analysis of protein yields across copolymer families** a. Coomassie-stained SDS–PAGE gels (ImageJ densitometry) and NanoTemper Panta Discovery fluorescence scans (330 nm) were used to quantify relative protein yields across six copolymer families (DIBMA, SMA, BZ, AASTY, Ultrasolute Amphipol, Cubipol). For each dataset, the number of values, minimum, maximum, percentiles (25%, median, 75%), mean, standard deviation, and 95% confidence intervals are reported. b. The Mann Whitney Test was used to test for a significant difference between SMA and the other copolymer classes. Both analyses demonstrate that next-generation copolymers (AASTY, Ultrasolute Amphipol, Cubipol) consistently display higher median values and narrower variance compared to classical SMA and DIBMA, underscoring their superior performance across targets.

**a**

| Target | Top1 | hdR (nm) | PDI | Top2 | hdR (nm) | PDI | Top 3 | hdR (nm) | PDI |
|---|---|---|---|---|---|---|---|---|---|
| HTR7 | Ultrasolute Amphipol 18 | 16.81 | 0.34 | Cubipol Glycerol | 17.6 | 0.21 | AASTY 11-50 | 18.08 | 0.27 |
| ADRB2 | Ultrasolute Amphipol 18 | 12.3 | 0.35 | AASTY 6-50 | 12.9 | 0.17 | Cubipol | 12.9 | 0.3 |
| AMYR3 (Calc+Ramp3) | Ultrasolute Amphipol 18 Biotin | 11.54 | 0.35 | Cubipol PEG | 11.56 | 0.31 | Ultrasolute Amphipol 18 | 11.57 | 0.33 |
| SLC1A6 | AASTY 6-50 | 10.03 | 0.2 | AASTY 11-50 | 10.18 | 0.23 | Glyco-Cubipol | 10.56 | 0.14 |
| SLC2A1 | SMA 200 | 6.89 | 0.31 | Ultrasolute Amphipol 18 | 8.31 | 0.44 | Cubipol | 9.49 | 0.23 |
| SLC15A4 | Cubipol Amin | 9.27 | 0.29 | Glyco-Cubipol | 9.44 | 0.48 | Cubipol Glycerol | 9.54 | 0.4 |
| hERG(t) | AASTY 6-50 | 10.28 | 0.24 | AASTY 11-50 | 10.35 | 0.28 | Glyco-Cubipol | 10.63 | 0.17 |
| CLDN1 | Glyco-DIBMA | 11.16 | 0.22 | SMA 200 | 11.18 | 0.22 | Cubipol Glycerol | 11.3 | 0.18 |
| CLDN3 | SMA 200 | 10.1 | 0.63 | BZ 25 | 10.6 | 0.62 | Cubipol Glycerol | 11.1 | 0.75 |
| CLDN4 | BZ 40 | 9.74 | 0.69 | Ultrasolute Amphipol 18 | 9.75 | 0.5 | Glyco-Cubipol | 9.82 | 0.52 |
| CD68 | Cubipol | 13.52 | 0.27 | BZ 40 | 13.59 | 0.51 | Sulfo-Cubipol | 13.87 | 0.25 |
| CD163 | Cubipol Glycerol | 16.43 | 0.25 | Sulfo-Cubipol | 16.92 | 0.24 | Glyco-DIBMA | 17.1 | 0.31 |
| EGFR | Cubipol Glycerol | 13.14 | 1.96 | SMA 200 | 14.49 | 0.17 | Ultrasolute Amphipol 18 | 14.53 | 0.3 |
| FAS | Ultrasolute Amphipol 18 | 10.17 | 0.41 | SMA 200 | 10.25 | 0.88 | AASTY 11-50 | 10.29 | 0.3 |

**b**

| Target | Copolymer | hdR (nm) | PDI |
|---|---|---|---|
| HTR7 | DIBMA-10 | 27.07 | 0.27 |
| | DIBMA Glycerol | 39.12 | 0.29 |
| | Glyco-DIBMA | 22.02 | 0.27 |
| | Sulfo-DIBMA | 35.97 | 0.28 |
| | SMA 200 | 21.33 | 0.24 |
| | BZ 25 | 19.72 | 0.26 |
| | BZ 40 | 23.45 | 0.2 |
| | Sulfo-SMA | 38.94 | 0.4 |
| | AASTY 6-50 | 20.42 | 0.62 |
| | AASTY 11-50 | 18.08 | 0.27 |
| | Ultrasolute Amphipol 18 | 16.81 | 0.34 |
| | Cubipol | 18.63 | 0.22 |
| | Sulfo-Cubipol | 22.04 | 0.22 |
| | Glyco-Cubipol | 18.41 | 0.21 |
| | Cubipol Glycerol | 17.6 | 0.21 |
| ADRB2 | DIBMA-10 | 17.6 | 0.35 |
| | DIBMA Glycerol | 20.1 | 0.5 |
| | Glyco-DIBMA | 16.8 | 0.77 |
| | Sulfo-DIBMA | 22.9 | 0.34 |
| | SMA 200 | 13.3 | 0.2 |
| | BZ 25 | 13.3 | 0.2 |
| | BZ 40 | 14.4 | 0.3 |
| | Sulfo-SMA | 13.1 | 0.19 |
| | AASTY 6-50 | 12.9 | 0.17 |
| | AASTY 11-50 | 34.3 | 0.23 |
| | Ultrasolute Amphipol 18 | 12.3 | 0.35 |
| | Cubipol | 12.9 | 0.3 |
| | Sulfo-Cubipol | 13.7 | 0.34 |
| | Glyco-Cubipol | 13 | 0.2 |
| | Cubipol Glycerol | 13.3 | 0.29 |
| AMYR3 (Calc+Ramp3) | DIBMA 6 | 54.12 | 1.23 |
| | AASTY 6-50 | 14.22 | 0.56 |
| | AASTY 11-50 | 12.28 | 0.32 |
| | Ultrasolute Amphipol 18 | 11.57 | 0.33 |
| | Cubipol | 14.15 | 0.43 |
| | Sulfo-Cubipol | 22.04 | 0.85 |
| | Glyco-Cubipol | 13.16 | 0.44 |
| | Cubipol Glycerol | 12.42 | 0.33 |
| | Cubipol PEG | 11.56 | 0.31 |
| | Cubipol Amin | 14.49 | 0.24 |
| | DIBMA 6 Biotin | 29.11 | 0.87 |
| | AASTY 11-50 Biotin | 12.81 | 0.26 |
| | Cubipol Biotin | 13.06 | 0.26 |
| | Sulfo-Cubipol Biotin | 44.17 | 1.22 |
| | Ultrasolute Amphipol 18 Biotin | 11.54 | 0.35 |
| SLC1A6 | DIBMA-10 | 15.65 | 0.36 |
| | DIBMA Glycerol | 19.51 | 0.46 |
| | Glyco-DIBMA | 11.12 | 0.24 |
| | Sulfo-DIBMA | 26.16 | 0.21 |
| | SMA 200 | 10.9 | 0.23 |
| | BZ 25 | 10.69 | 0.18 |
| | BZ 40 | 11.76 | 0.15 |
| | Sulfo-SMA | 30.82 | 0.13 |
| | AASTY 6-50 | 10.03 | 0.2 |
| | AASTY 11-50 | 10.18 | 0.23 |
| | Ultrasolute Amphipol 18 | 10.64 | 0.13 |
| | Cubipol | 11.14 | 0.23 |
| | Sulfo-Cubipol | 11.9 | 0.27 |
| | Glyco-Cubipol | 10.56 | 0.14 |
| | Cubipol Glycerol | 10.79 | 0.18 |
| SLC2A1 | DIBMA-10 | 16.98 | 0.16 |
| | DIBMA Glycerol | 18.41 | 0.19 |
| | Glyco-DIBMA | 10.14 | 0.23 |
| | Sulfo-DIBMA | 24.51 | 0.94 |
| | SMA 200 | 6.89 | 0.31 |
| | BZ 25 | 16.95 | 0.14 |
| | BZ 40 | 19.65 | 0.2 |
| | Sulfo-SMA | 27.12 | 0.13 |
| | AASTY 6-50 | 29406.83 | 2.22 |
| | AASTY 11-50 | 15.34 | 0.19 |
| | Ultrasolute Amphipol 18 | 8.31 | 0.44 |
| | Cubipol | 9.49 | 0.23 |
| | Sulfo-Cubipol | 15.34 | 0.2 |
| | Glyco-Cubipol | 10.41 | 0.34 |
| | Cubipol Glycerol | 9.52 | 0.26 |
| SLC15A4 | DIBMA 6 | 18.4 | 0.42 |
| | AASTY 6-50 | 12.62 | 0.19 |
| | AASTY 11-50 | 11.53 | 0.77 |
| | Ultrasolute Amphipol 18 | 10.25 | 0.61 |
| | Cubipol | 9.74 | 0.35 |
| | Sulfo-Cubipol | 19.25 | 0.41 |
| | Glyco-Cubipol | 9.44 | 0.48 |
| | Cubipol Glycerol | 9.54 | 0.4 |
| | Cubipol PEG | 10.28 | 0.46 |
| | Cubipol Amin | 9.27 | 0.29 |
| | DIBMA 6 Biotin | 23.01 | 0.54 |
| | AASTY 11-50 Biotin | 11.17 | 0.41 |
| | Cubipol Biotin | 10.1 | 0.53 |
| | Sulfo-Cubipol Biotin | 23.2 | 0.27 |
| | Ultrasolute Amphipol 18 Biotin | 11.45 | 0.52 |
| hERG(t) | DIBMA-10 | 12.54 | 0.21 |
| | DIBMA Glycerol | 17.28 | 0.45 |
| | Glyco-DIBMA | 13.05 | 0.17 |
| | Sulfo-DIBMA | 40.65 | 0.38 |
| | SMA 200 | 10.69 | 0.27 |
| | BZ 25 | 13.28 | 0.35 |
| | BZ 40 | 11.97 | 0.2 |
| | Sulfo-SMA | 35.31 | 0.22 |
| | AASTY 6-50 | 10.28 | 0.24 |
| | AASTY 11-50 | 10.35 | 0.28 |
| | Ultrasolute Amphipol 18 | 10.95 | 0.57 |
| | Cubipol | 11.33 | 0.24 |
| | Sulfo-Cubipol | 13.31 | 0.3 |
| | Glyco-Cubipol | 10.63 | 0.17 |
| | Cubipol Glycerol | 11.13 | 0.15 |
| CLDN1 | DIBMA-10 | 13.11 | 0.31 |
| | DIBMA Glycerol | 14.46 | 0.39 |
| | Glyco-DIBMA | 11.16 | 0.22 |
| | Sulfo-DIBMA | 23.27 | 0.18 |
| | SMA 200 | 11.18 | 0.22 |
| | BZ 25 | 11.47 | 0.34 |
| | BZ 40 | 11.78 | 0.18 |
| | Sulfo-SMA | 30.82 | 0.21 |
| | AASTY 6-50 | 12.64 | 0.2 |
| | AASTY 11-50 | 12.08 | 0.19 |
| | Ultrasolute Amphipol 18 | 12.17 | 0.29 |
| | Cubipol | 11.86 | 0.31 |
| | Sulfo-Cubipol | 18.39 | 0.29 |
| | Glyco-Cubipol | 11.85 | 0.23 |
| | Cubipol Glycerol | 11.3 | 0.18 |
| CLDN3 | DIBMA-10 | 25.1 | 0.97 |
| | DIBMA Glycerol | 16.8 | 0.49 |
| | Glyco-DIBMA | 547.1 | 1.56 |
| | Sulfo-DIBMA | 26.8 | 0.73 |
| | SMA 200 | 10.1 | 0.63 |
| | BZ 25 | 10.6 | 0.62 |
| | BZ 40 | 17.2 | 1.01 |
| | Sulfo-SMA | 24.1 | 0.83 |
| | AASTY 6-50 | 12.6 | 0.71 |
| | AASTY 11-50 | 13.3 | 0.58 |
| | Ultrasolute Amphipol 18 | 13.5 | 0.59 |
| | Cubipol | 41.6 | 1.36 |
| | Sulfo-Cubipol | 23.7 | 0.85 |
| | Glyco-Cubipol | 15 | 0.95 |
| | Cubipol Glycerol | 11.1 | 0.75 |
| CLDN4 | DIBMA-10 | 11.2 | 0.55 |
| | DIBMA Glycerol | 14.2 | 0.79 |
| | Glyco-DIBMA | 17.4 | 1.05 |
| | Sulfo-DIBMA | 19.1 | 0.3 |
| | SMA 200 | 2847.9 | 1.1 |
| | BZ 25 | 9.86 | 0.84 |
| | BZ 40 | 9.74 | 0.69 |
| | Sulfo-SMA | 16.7 | 0.62 |
| | AASTY 6-50 | 13.5 | 0.94 |
| | AASTY 11-50 | 17.6 | 1.05 |
| | Ultrasolute Amphipol 18 | 9.75 | 0.5 |
| | Cubipol | 52.3 | 1.48 |
| | Sulfo-Cubipol | 21.8 | 0.8 |
| | Glyco-Cubipol | 9.82 | 0.52 |
| | Cubipol Glycerol | 93639.5 | 1.97 |
| CD68 | DIBMA-10 | 16.64 | 0.28 |
| | DIBMA Glycerol | 16.31 | 0.38 |
| | Glyco-DIBMA | 14.52 | 0.37 |
| | Sulfo-DIBMA | 22.44 | 0.45 |
| | SMA 200 | 14.05 | 0.32 |
| | BZ 25 | 14.91 | 0.32 |
| | BZ 40 | 13.59 | 0.51 |
| | Sulfo-SMA | 35.65 | 0.18 |
| | AASTY 6-50 | 15.37 | 0.29 |
| | AASTY 11-50 | 14.01 | 0.24 |
| | Ultrasolute Amphipol 18 | 14.17 | 0.19 |
| | Cubipol | 13.52 | 0.27 |
| | Sulfo-Cubipol | 13.87 | 0.25 |
| | Glyco-Cubipol | 14.04 | 0.23 |
| | Cubipol Glycerol | 14.78 | 0.15 |
| CD163 | DIBMA-10 | 25.38 | 0.33 |
| | DIBMA Glycerol | 47714.72 | 1.04 |
| | Glyco-DIBMA | 17.1 | 0.31 |
| | Sulfo-DIBMA | 28.45 | 0.61 |
| | SMA 200 | 17.92 | 0.32 |
| | BZ 25 | 18.85 | 0.32 |
| | BZ 40 | 20.15 | 0.33 |
| | Sulfo-SMA | 36.63 | 0.27 |
| | AASTY 6-50 | 17.77 | 0.41 |
| | AASTY 11-50 | 17.26 | 0.35 |
| | Ultrasolute Amphipol 18 | 17.68 | 0.81 |
| | Cubipol | 26.96 | 1.04 |
| | Sulfo-Cubipol | 16.92 | 0.24 |
| | Glyco-Cubipol | 17.58 | 0.58 |
| | Cubipol Glycerol | 16.43 | 0.25 |
| EGFR | DIBMA-10 | 17.35 | 0.31 |
| | DIBMA Glycerol | 19.84 | 0.42 |
| | Glyco-DIBMA | 15.41 | 0.24 |
| | Sulfo-DIBMA | 20.55 | 0.39 |
| | SMA 200 | 14.49 | 0.17 |
| | BZ 25 | 15.18 | 0.22 |
| | BZ 40 | 16.25 | 0.18 |
| | Sulfo-SMA | 30.76 | 0.53 |
| | AASTY 6-50 | 14.91 | 0.19 |
| | AASTY 11-50 | 14.98 | 0.28 |
| | Ultrasolute Amphipol 18 | 14.53 | 0.3 |
| | Cubipol | 15.21 | 0.19 |
| | Sulfo-Cubipol | 15.22 | 0.23 |
| | Glyco-Cubipol | 14.9 | 0.21 |
| | Cubipol Glycerol | 13.14 | 1.96 |
| FAS | DIBMA-10 | 13.8 | 0.36 |
| | DIBMA Glycerol | 13.1 | 0.24 |
| | Glyco-DIBMA | 11.17 | 0.3 |
| | Sulfo-DIBMA | 33.01 | 0.13 |
| | SMA 200 | 10.25 | 0.88 |
| | BZ 25 | 13.6 | 0.6 |
| | BZ 40 | 11.4 | 0.48 |
| | Sulfo-SMA | 37.6 | 0.15 |
| | AASTY 6-50 | 10.57 | 0.26 |
| | AASTY 11-50 | 10.29 | 0.3 |
| | Ultrasolute Amphipol 18 | 10.17 | 0.41 |
| | Cubipol | 10.72 | 0.3 |
| | Sulfo-Cubipol | 14.02 | 0.5 |
| | Glyco-Cubipol | 11.17 | 0.3 |
| | Cubipol Glycerol | 12.65 | 0.4 |

**Extended Data Figure 5. Dynamic light scattering analysis of 14 membrane protein targets.** a. Top-3 performing copolymers for each of the 14 tested membrane proteins, ranked by the combination of smallest hydrodynamic radius (hdR) and lowest polydispersity index (PDI). b. Complete hdR and

PDI dataset for all copolymer-target combinations across the 14 proteins, providing a comprehensive overview of nanodisc size and homogeneity.

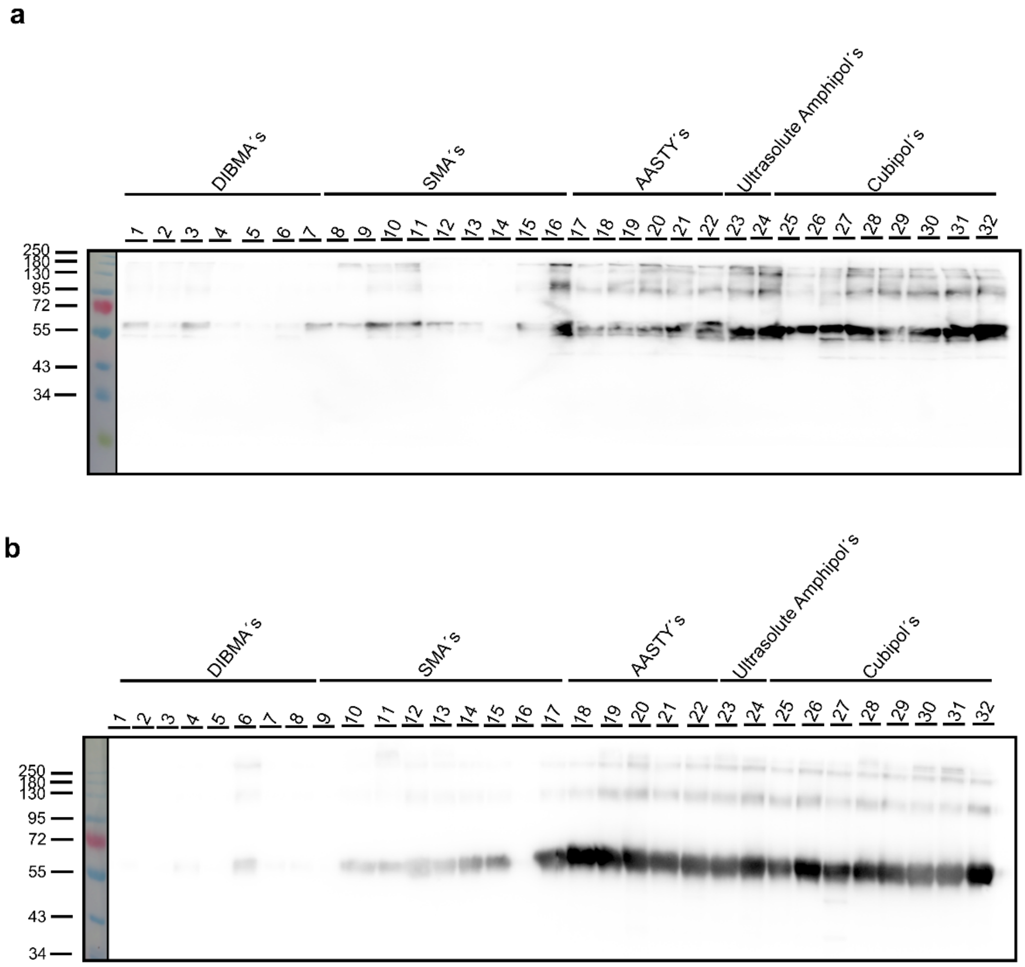


**Extended Data Figure 6. Western blot validation of P2X4 copolymer screens.** Western blots corresponding to the full copolymer screens of P2X4 expressed in *Tni* insect cells and HEK293 mammalian cells. The blots confirm protein identity and recovery across 32 different copolymer conditions.

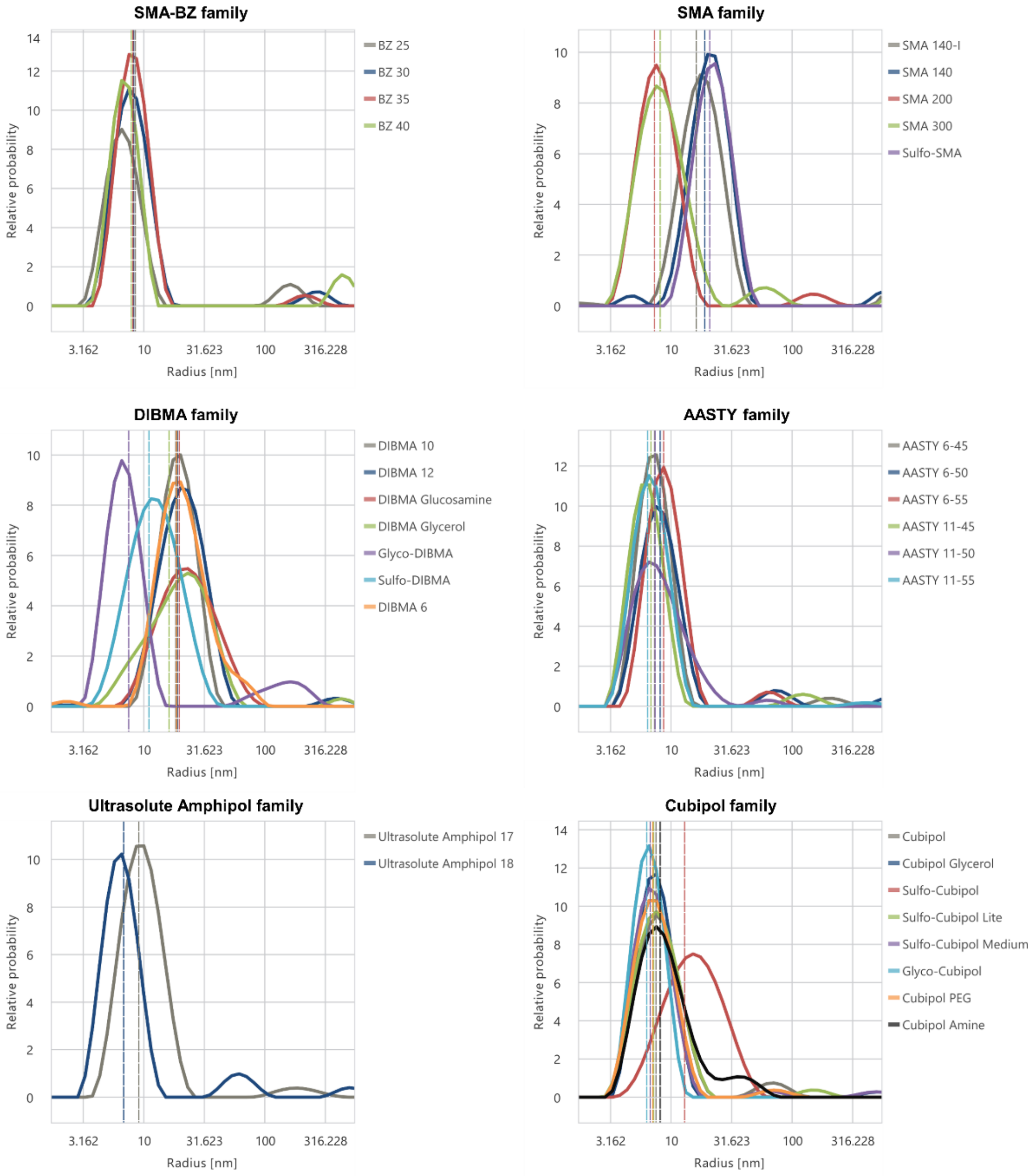


**Extended Data Figure 7. Dynamic light scattering (DLS) analysis of P2X4$^{HEK}$ in 32 copolymers.**

Full DLS dataset of P2X4 expressed in HEK293 cells and solubilized with the complete NativeMP™ copolymer suite, grouped by backbone families. For each condition, hydrodynamic radius (hdR) and polydispersity index (PDI) were determined to assess nanodisc size and homogeneity. In Fig. 4b, only three representative examples (SMA200, AASTY 6-55, Glyco-Cubipol) are shown; here the complete set is presented for reference. These values provide the basis for the composite scoring and ranking shown in Fig. 4e.

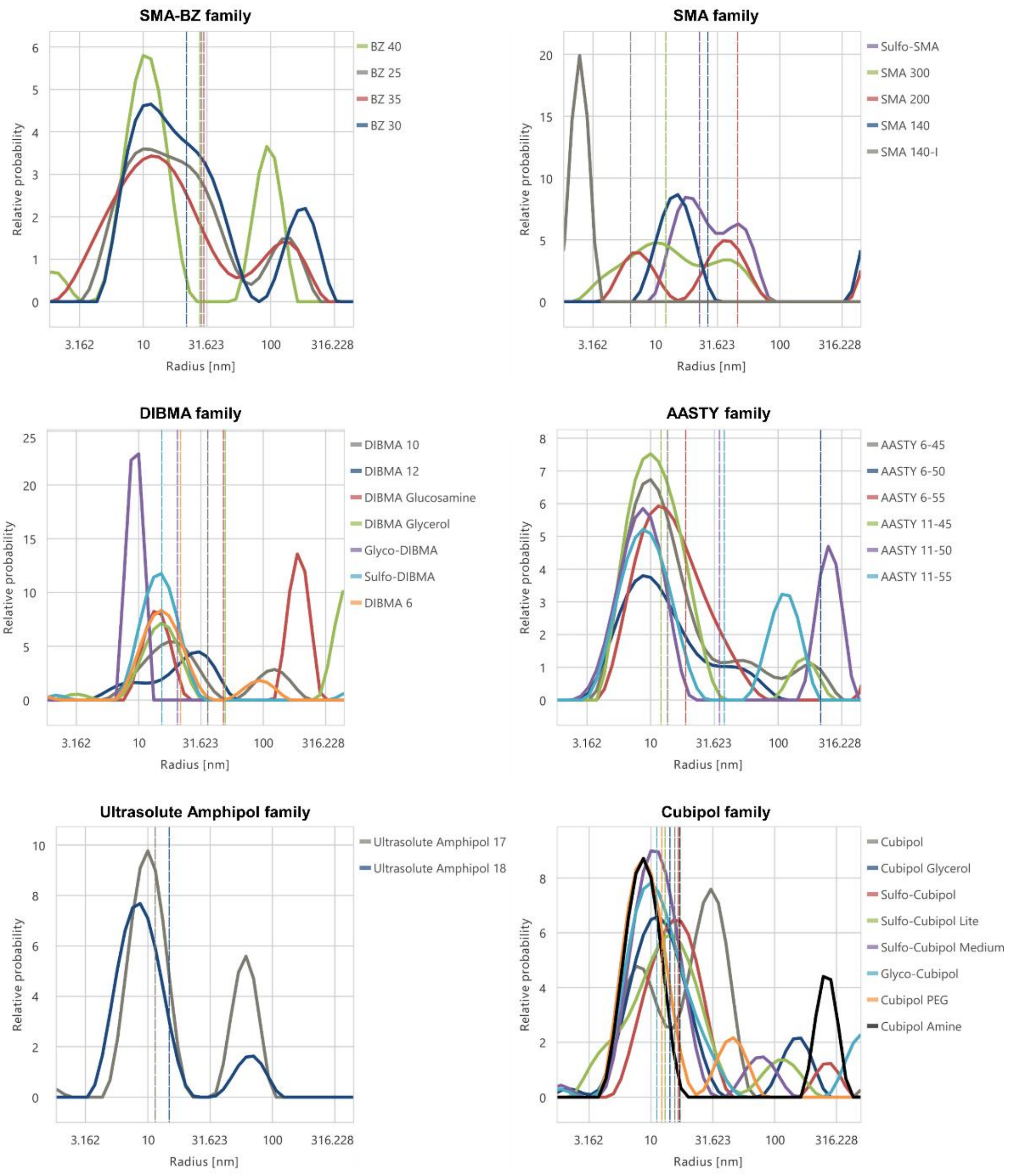


**Extended Data Figure 8. Dynamic light scattering (DLS) analysis of P2X4$^{Tni}$ in 32 copolymers.**

Full DLS dataset of P2X4 expressed in *T.ni* insect cells and solubilized with the complete NativeMP™ copolymer suite, grouped by backbone families. Hydrodynamic radius (hdR) and polydispersity index (PDI) values are displayed for all copolymer conditions. This analysis mirrors the approach in Fig. 4b and Supplementary Figure S9 but highlights the generally higher heterogeneity and reduced stability observed for the *T.ni*-derived construct. As for the HEK-derived protein, these data directly underlie the scoring and comparative ranking presented in Fig. 4e.

| Rank | Copolymer | Tm_IP mean | Tm_IP stand dev | Tm_IP mean purification 1 | Tm_IP mean purification 2 | hDR mean | hDR stand dev | hDR purification 1 | hDR purification 2 | PDI mean | Turbidity mean | Elution Quantity | Tm_IP norm | hdR norm | PDI norm | Turbidity norm | Elution Quantity norm | Score |
|---|---|---|---|---|---|---|---|---|---|---|---|---|---|---|---|---|---|---|
| 1 | Glyco-Cubipol | 64,61 | 0,39 | 65,00 | 64,22 | 7,20 | 0,85 | 6,35 | 8,05 | 0,14 | 45,93 | 91,85 | 0,87 | 1,00 | 1,00 | 0,76 | 0,92 | 0,92 |
| 2 | Sulfo-Cubipol Medium | 64,47 | 2,32 | 66,79 | 62,15 | 7,69 | 0,91 | 6,79 | 8,60 | 0,18 | 59,01 | 87,31 | 0,96 | 1,00 | 0,92 | 0,50 | 0,87 | 0,92 |
| 3 | Cubipol PEG | 63,81 | 0,73 | 64,54 | 63,08 | 8,09 | 1,04 | 7,05 | 9,13 | 0,25 | 47,52 | 100,00 | 0,85 | 1,00 | 0,80 | 0,73 | 1,00 | 0,91 |
| 4 | Cubipol Amine | 66,32 | 1,27 | 67,59 | 65,05 | 8,53 | 0,36 | 8,17 | 8,89 | 0,29 | 85,02 | 83,93 | 1,00 | 1,00 | 0,71 | 0,00 | 0,84 | 0,88 |
| 5 | Cubipol Glycerol | 63,90 | 0,57 | 64,47 | 63,33 | 7,86 | 0,77 | 7,10 | 8,63 | 0,19 | 67,09 | 85,06 | 0,84 | 1,00 | 0,90 | 0,35 | 0,85 | 0,87 |
| 6 | Cubipol | 63,16 | 0,17 | 63,33 | 62,98 | 9,07 | 1,51 | 7,56 | 10,58 | 0,22 | 41,03 | 83,29 | 0,79 | 1,00 | 0,85 | 0,85 | 0,83 | 0,86 |
| 7 | Ultrasolute Amphipol 18 | 62,23 | 0,08 | 62,15 | 62,31 | 7,68 | 0,82 | 6,86 | 8,50 | 0,41 | 58,99 | 95,93 | 0,73 | 1,00 | 0,49 | 0,50 | 0,96 | 0,82 |
| 8 | Ultrasolute Amphipol 17 | 61,19 | 0,03 | 61,22 | 61,16 | 8,70 | 0,41 | 9,12 | 8,29 | 0,18 | 63,68 | 79,42 | 0,68 | 1,00 | 0,92 | 0,41 | 0,79 | 0,80 |
| 9 | AASTY 11-45 | 58,38 | 0,07 | 58,31 | 58,45 | 7,80 | 0,99 | 6,81 | 8,79 | 0,44 | 41,99 | 101,25 | 0,54 | 1,00 | 0,43 | 0,83 | 1,01 | 0,78 |
| 10 | AASTY 11-55 | 57,20 | 0,78 | 57,98 | 56,42 | 8,45 | 1,97 | 6,47 | 10,42 | 0,17 | 47,70 | 80,52 | 0,52 | 1,00 | 0,94 | 0,72 | 0,81 | 0,76 |
| 11 | AASTY 6-50 | 59,65 | 0,92 | 60,57 | 58,73 | 8,78 | 0,56 | 8,22 | 9,33 | 0,37 | 39,73 | 70,81 | 0,65 | 1,00 | 0,56 | 0,88 | 0,71 | 0,75 |
| 12 | AASTY 11-50 | 58,84 | 0,38 | 58,46 | 59,22 | 7,81 | 0,33 | 7,48 | 8,14 | 0,33 | 78,59 | 88,42 | 0,54 | 1,00 | 0,64 | 0,12 | 0,88 | 0,73 |
| 13 | AASTY 6-45 | 57,62 | 0,81 | 58,42 | 56,81 | 8,11 | 0,83 | 7,28 | 8,94 | 0,22 | 63,23 | 53,40 | 0,54 | 1,00 | 0,85 | 0,42 | 0,53 | 0,68 |
| 14 | Sulfo-DIBMA | 63,68 | 1,54 | 65,22 | 62,14 | 11,92 | 0,92 | 11,00 | 12,84 | 0,25 | 66,04 | 14,91 | 0,88 | 0,93 | 0,80 | 0,37 | 0,15 | 0,68 |
| 15 | Sulfo-Cubipol | 59,28 | 0,99 | 60,27 | 58,29 | 13,44 | 0,35 | 13,10 | 13,79 | 0,26 | 34,09 | 49,10 | 0,63 | 0,79 | 0,77 | 0,99 | 0,49 | 0,67 |
| 16 | BZ 30 | 58,37 | 0,37 | 58,74 | 58,01 | 9,48 | 1,15 | 8,33 | 10,63 | 0,40 | 53,17 | 46,69 | 0,56 | 1,00 | 0,50 | 0,62 | 0,47 | 0,64 |
| 17 | BZ 25 | 59,13 | 0,15 | 59,29 | 58,98 | 9,80 | 1,21 | 8,59 | 11,02 | 0,67 | 69,74 | 66,26 | 0,58 | 1,00 | 0,00 | 0,30 | 0,66 | 0,64 |
| 18 | Glyco-DIBMA | 66,71 | 0,00 | 66,71 |  | 13,29 | 5,69 | 7,60 | 18,98 | 0,48 | 84,34 | 3,95 | 0,96 | 1,00 | 0,36 | 0,01 | 0,04 | 0,63 |
| 19 | BZ 40 | 57,46 | 0,71 | 58,18 | 56,75 | 9,16 | 1,29 | 7,87 | 10,45 | 0,43 | 41,90 | 23,53 | 0,53 | 1,00 | 0,45 | 0,84 | 0,24 | 0,58 |
| 20 | Sulfo-Cubipol Lite | 55,80 | 8,21 | 47,59 | 64,01 | 8,31 | 1,11 | 7,20 | 9,41 | 0,19 | 74,36 | 75,84 | 0,00 | 1,00 | 0,91 | 0,21 | 0,76 | 0,54 |
| 21 | SMA 300 | 50,68 | 0,74 | 51,42 | 49,94 | 9,82 | 1,58 | 8,24 | 11,40 | 0,37 | 46,87 | 40,09 | 0,19 | 1,00 | 0,56 | 0,74 | 0,40 | 0,51 |
| 22 | SMA 200 | 50,96 | 0,00 | 50,96 |  | 8,01 | 0,73 | 7,28 | 8,74 | 0,28 | 33,45 | 27,25 | 0,17 | 1,00 | 0,74 | 1,00 | 0,27 | 0,50 |
| 23 | DIBMA 6 | 58,34 | 0,00 | 58,34 |  | 20,43 | 1,52 | 18,92 | 21,95 | 0,21 | 39,17 | 4,82 | 0,54 | 0,41 | 0,87 | 0,89 | 0,05 | 0,43 |
| 24 | DIBMA 10 | 55,30 | 0,00 | 55,30 |  | 19,29 | 0,69 | 18,59 | 19,98 | 0,33 | 69,57 | 2,05 | 0,39 | 0,43 | 0,65 | 0,30 | 0,02 | 0,33 |
| 25 | DIBMA 12 | 56,01 | 0,00 | 56,01 |  | 49,10 | 30,04 | 19,05 | 79,14 | 0,33 | 82,03 | 2,33 | 0,42 | 0,40 | 0,64 | 0,06 | 0,02 | 0,32 |
| 26 | BZ 35 | 59,33 | 0,61 | 59,94 | 58,72 | 9,43 | 1,20 | 8,23 | 10,63 | 0,25 |  | 44,44 | 0,62 | 1,00 | 0,78 |  | 0,44 |  |
| 27 | AASTY 6-55 | 43,53 | 13,21 | 56,74 | 30,32 | 8,89 | 0,20 | 8,69 | 9,08 | 0,25 |  | 85,97 | 0,46 | 1,00 | 0,80 |  | 0,86 |  |

**Extended Data Figure 9. Comprehensive ranking table of copolymer screening for P2X4 full-length in HEK293 cells.** Top 27 of all 32 tested copolymers are listed with quantitative values from two independent purifications. Parameters include melting temperature ($T_{mIP}$ ), hydrodynamic radius (hdR), polydispersity index (PDI), turbidity, and elution quantity. Mean values, standard deviations, and results from both purifications are reported. Normalized values for each parameter were calculated and integrated into a composite score (max = 1) using predefined weights (Tm 0.35, radius 0.25, PDI 0.10, turbidity 0.05, elution quantity 0.25). The final score was used to generate a rank order (1 = best). Top-performing copolymers were predominantly Cubipol variants (Glyco-Cubipol, Sulfo-Cubipol Medium, Cubipol PEG, Cubipol Amine, and Cubipol Glycerol), while several DIBMA and SMALP derivatives occupied lower ranks due to larger particle sizes, higher PDI, or reduced recovery.

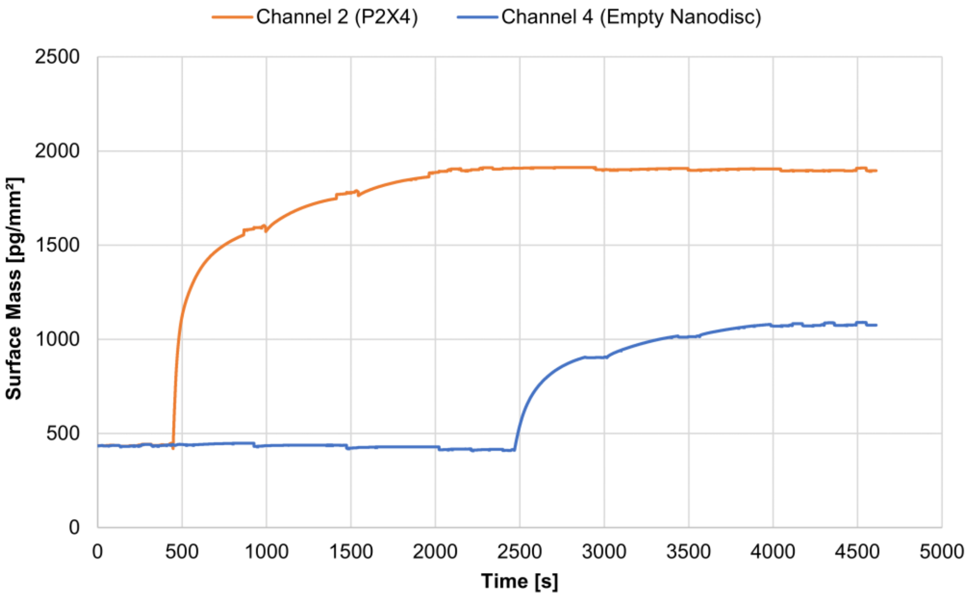


**Extended Data Figure 10. Immobilization of biotinylated nanodiscs on streptavidin chips.**

Biotinylated U18 nanodiscs containing P2X4 (Protein nanodisc) and control copolymer-host lipid nanodisc (Empty Nanodisc) were immobilized on streptavidin-coated chips. The immobilization levels reached ~1470 pg $mm^{-2}$ for Protein nanodisc on channel 2 and ~639 pg $mm^{-2}$ for Empty Nanodisc on channel 4, demonstrating efficient coupling.

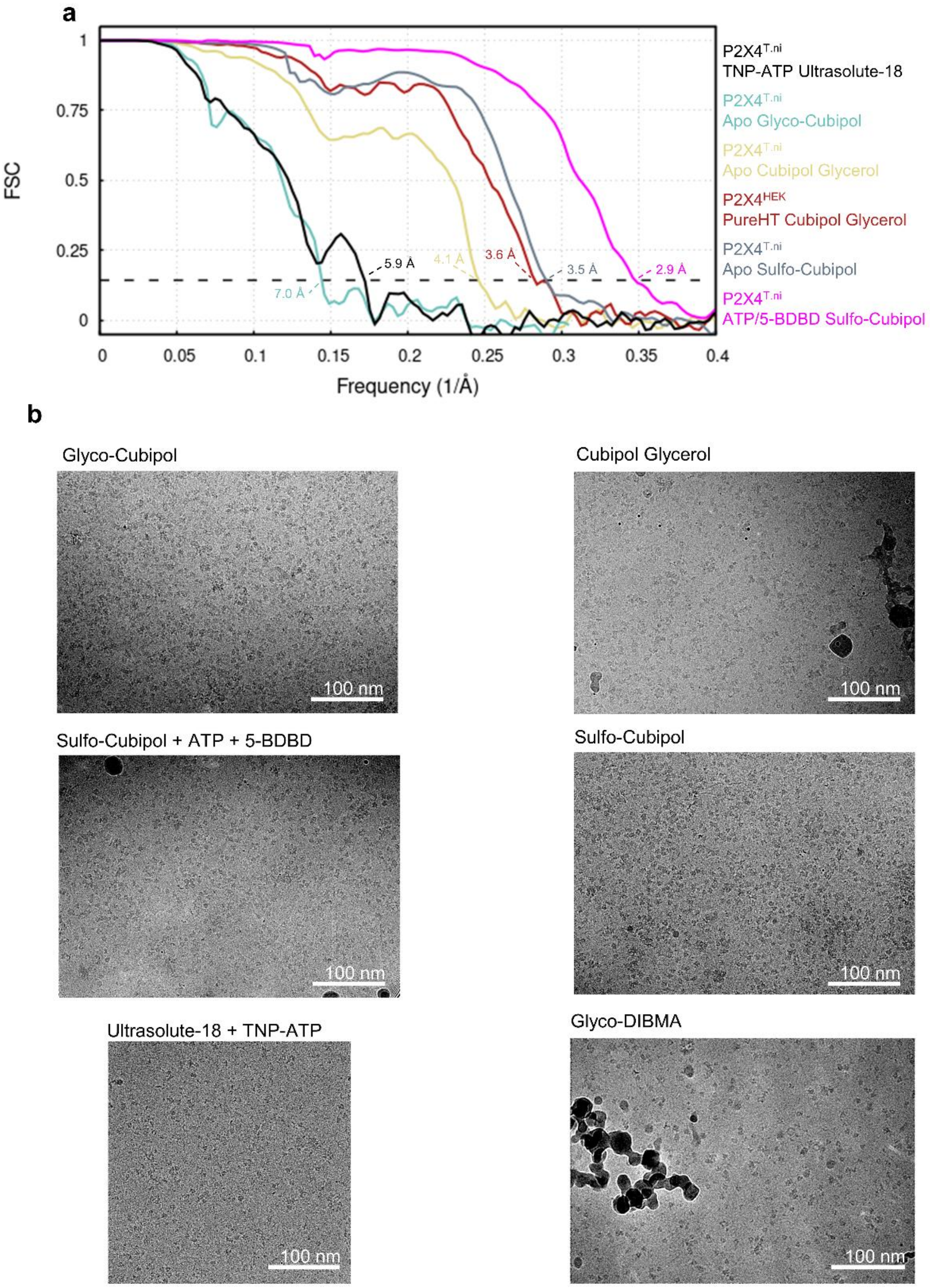


**Extended Data Figure 11. Resolution and representative images of the Cryo-EM screening.**

**a**. Fourier-shell-correlation estimation of the resolution of all reconstructed maps shown in the article. In all cases, we used a threshold of 0.143, which is highlighted by a segmented line in the plot. **b**. Representative micrographs of the different polymers tested for P2X4$^{T.ni}$ in Cryo-EM.

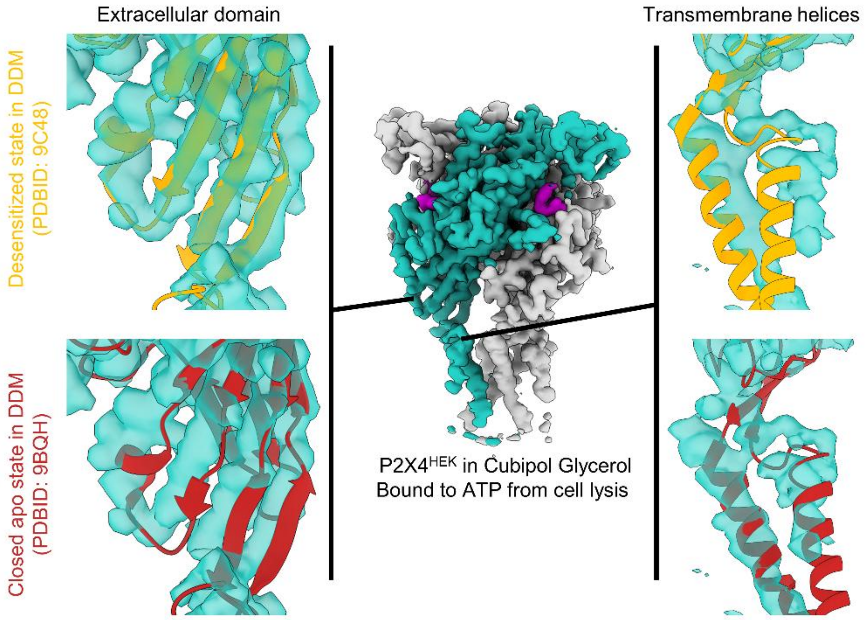


**Extended Data Figure 12**. **Comparison of the conformational state of P2X4$^{HEK}$ in Cubipol Glycerol purified without automated system with the available structures of P2X4 in detergent.**

The structures of the desensitized (PDBID: 9C48) and closed (PDBID: 9BQH) states were rigid-body fitted into our Cryo-EM map. Consistent with the presence of ATP, the conformation of the extracellular domain fits the desensitized state. The conformation of the transmembrane helices however, does not fit well either state, suggesting that P2X4$^{HEK}$ is in a different conformation in our samples.

**Table S1.** Summary of Cryo-EM data collection strategy

| Sample | Glycerol-Cubipol (P2X4$^{HEK}$) | Glyco-DIBMA (P2X4$^{T.ni}$) | Ultrasolute-18 (P2X4$^{T.ni}$) | Glyco-Cubipol (P2X4$^{T.ni}$) | Glycerol-Cubipol (P2X4$^{T.ni}$) | Sulfo-Cubipol (P2X4$^{T.ni}$) | Sulfo-Cubipol (P2X4$^{T.ni}$) |
|---|---|---|---|---|---|---|---|
| **Accession codes** | EMD-59335 | | EMD-59364 | EMD-59363 | EMD-59361 | EMD-59333 pdb_000033BX | EMD-59354 pdb_000033CQ |
| **Ligands** | | | 100 µM TNP-ATP | | | | 100 µM ATP 100 µM 5-BDBD |
| **Microscope** | Talos Arctica | | Titan Krios G4 | | | | |
| **Acceleration voltage (kV)** | 200 | | 300 | | | | |
| **Total Dose (e-/A2)** | 65 | 65 | 65 | 60 | 60 | 65 | 65 |
| **Exposure time (s)** | 3.6 | 3.6 | 2.6 | 2.25 | 2.25 | 2.6 | 2.6 |
| **Camera** | K3 | | Falcon 4i | K3 | | | |
| **Energy filter** | Gatan BioQuantum (20 eV) | | - | Gatan BioContinuum (20 eV) | | | |
| **Pixel size (Å)** | 0.816 | 0.816 | 0.83 | 0.82 | 0.82 | 0.82 | 0.82 |
| **Number of micrographs** | 1,400 | 2,194 | 5,061 | 754 | 1,098 | 3,195 | 3,619 |
| **Number of picked particles** | 496,656 | 168,863 | 291,171 | 119,143 | 179,782 | 543,282 | 752,494 |
| **Number of particles in reconstruction** | 13,649 | - | 23,647 | 15,709 | 12,375 | 39,110 | 43,330 |
| **Resolution (Å)** | 3.6[a] | - | 5.9 | 7.0 | 4.1[a] | 3.5[a] | 2.9[a] |

[a]Resolution calculated after Bayesian polishing and CTF refinement

### Table S2. Atomic model refinement statistics

| | **P2X4$^{T.ni}$ Sulfo-Cubipol** (EMDB-59333) (PDB 33BX) | **P2X4$^{T.ni}$/ATP Sulfo-Cubipol** (EMDB-59354) (PDB 33CQ) |
|---|---|---|
| **Model Refinement** | | |
| Initial model used | ModelAngelo | ModelAngelo |
| Model composition | | |
| Non-hydrogen atoms | 7578 | 8217 |
| Protein residues | 942 | 993 |
| Ligands | - | 3 ATP/3 Mg |
| *B* factors (Å$^2$) | | |
| Protein | 130.24 | 125.4 |
| Ligand | - | 69.2 |
| R.m.s. deviations | | |
| Bond lengths (Å) | 0.0129 | 0.0135 |
| Bond angles (°) | 2.04 | 1.94 |
| Validation | | |
| MolProbity score | 1.45 | 1.62 |
| Clashscore | 3.72 | 3.99 |
| Poor rotamers (%) | 0.0 | 0.0 |
| Ramachandran plot | | |
| Favored (%) | 95.81 | 93.31 |
| Allowed (%) | 4.19 | 6.69 |
| Disallowed (%) | 0.0 | 0.0 |
